# Neural signatures of temporal anticipation in human cortex represent event probability density

**DOI:** 10.1101/2024.11.20.624477

**Authors:** Matthias Grabenhorst, David Poeppel, Georgios Michalareas

## Abstract

Temporal prediction is a fundamental function of neural systems. Recent advances suggest that humans anticipate future events by calculating probability density functions, rather than hazard rates. However, direct neural evidence for this mechanism is lacking. We recorded neural activity using magnetoencephalography as participants anticipated auditory and visual events distributed in time. We show that temporal anticipation, measured as reaction times, approximates the event probability density function, but not hazard rate. Temporal anticipation manifests as spatiotemporally patterned activity in three anatomically and functionally distinct parieto-temporal and sensorimotor cortical areas. In both audition and vision, each of these areas revealed a marked neural signature of anticipation: Prior to sensory cues, activity in a specific frequency band of neural oscillations, spanning alpha and beta ranges, encodes the event probability density function. Strikingly, these neural signals predicted reaction times to imminent sensory cues. These results show that supra-modal representations of probability density across cortex underlie the anticipation of future events.

## Introduction

The anticipation of future events is foundational to many complex functions, such as associative learning^1-3^, decision-making^4,5^, and motor-preparation^6,7^. For example, a predator predicts its prey’s movements, decides to quickly attack, and secures the next meal. A human boxer anticipates her opponent’s actions, moves rapidly, and evades the attack. In communication, humans and animals track, predict, and quickly react to their partners’ auditory and visual signals. Accordingly, the temporal prediction of sensory cues is considered an elementary function of neural systems^8-10^.

Temporal prediction requires the estimation of elapsed time relative to a reference point and the estimation of event probability over time. Understanding the brain mechanisms that solve this computational problem has been an important goal in neuroscience^11-18^. Still, the cortical representation of temporal event statistics is not understood.

Neural signals reflecting basic temporal prediction are observed across the cortical hierarchy. These signals are often associated with enhanced sensory perception and improved motor performance. In rodent primary auditory cortex^9^ and non-human primate’s early (V1) ^19^ and late (V4) ^20^ visual areas, as well as premotor cortex^5^, neuronal firing rates reflect the expectation of events in time. Likewise, in humans, expectations modulate activity in auditory^21^ and visual^22^ cortices. Specifically, spectral power modulations in the alpha (8–13 Hz) ^23^ and beta (15–30 Hz) ^17^ bands are related to prediction in (sensori-)motor activity.

It is debated how such expectation signals emerge in sensory and motor regions^4,12^. There is evidence that prediction may, at least partially, be an intrinsic property of sensory systems^24,25^ that, in addition to top-down modulatory effects^26^, shapes perception. Motor cortex function, on the other hand, critically depends on sensorimotor transformation^27^.

The posterior parietal cortex (PPC) is essential for motor preparation, based on its connectivity to both sensory and action-planning systems^28,29^. An influential hypothesis of temporal prediction proposes a specific computation in PPC based on the hazard rate (HR) of events^11,30,31^. In words, the HR represents the probability that a future event is imminent, given that it has not yet occurred^30^. The HR is widely assumed to be the canonical computation underlying temporal anticipation^3,11-17,19,20,32-35^. Specifically, the HR is argued to be represented in lateral intraparietal area LIP in non-human primates, potentially driving eye saccades to anticipated future events^11^. In humans, the HR computation is associated with several cortical areas, including early visual cortex and inferior parietal lobule^32^, and motor cortex^17,32^. However, the computation of HR is a complex^36^ and numerically unstable^37^ mathematical operation which questions its neurobiological plausibility and raises doubts about the HR’s generality as a neural prediction signal. This highlights a long-standing question: What is the cortical representation of event probability across time?

The probability density function of events over time, the *event PDF*, is a computationally simpler variable than the HR. The event PDF can be empirically approximated by two basic operations: the estimation of elapsed time and the counting of events over time, as more and more events are registered (Fig. 1a). Recently, a model of anticipatory behavior based on the event PDF was proposed^38,39^.

**Fig. 1.**
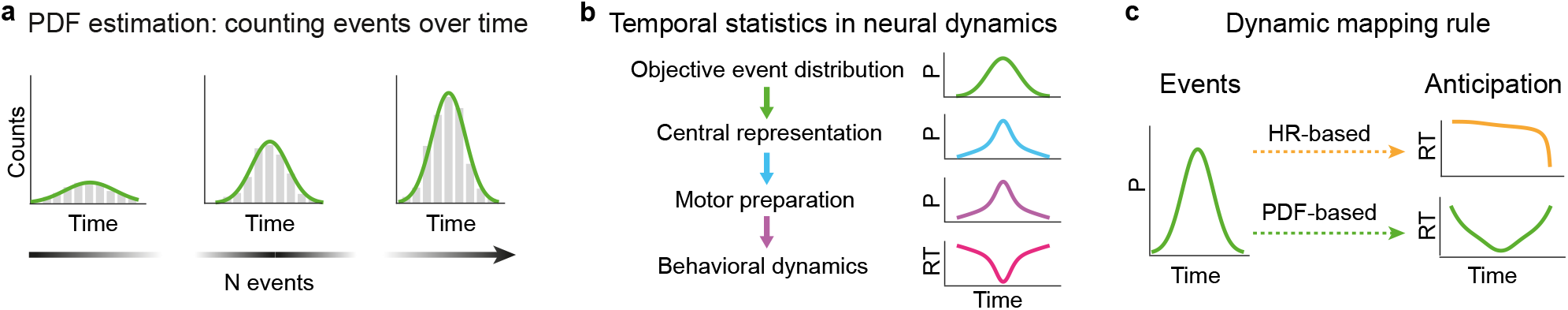
Conceptual hypotheses in temporal anticipation. **a**. Neural systems estimate the event probability density function (PDF) by counting events over time. **b**. A central representation of temporal statistics, e.g. probability over time, shapes motor preparation leading to adaptive behavior: where event probability is high, reaction time is short and vice versa. **c**. Temporal anticipation as a computational mapping problem. To adjudicate between computational hypotheses, the objective event PDF is related to reaction time, a proxy for anticipation, by different mapping rules (hazard-rate-based and PDF-based, **Methods**).

The current study aimed to identify the cortical representation of temporal statistics by investigating HR-based and PDF-based models of anticipation to adjudicate between them. We are guided by two general hypotheses: i) Inference of the temporal statistics of sensory events leads to a neural representation of probability over time. ii) This representation of sensory statistics ultimately shapes the preparation and execution of a fast motor response, the behavioral hallmark of successful prediction^30,40^. Specifically, we assume an inverse relationship between event probability and motor preparation: when event probability is high, reaction time should be short and vice versa (Fig. 1b).

We conceptualize temporal anticipation as a computational mapping problem where a neural representation of temporal statistics relates stochastic sensory input to anticipatory behavior, expressed as reaction time (RT) modulation (Fig. 1c). We performed experiments in audition and vision, two modalities that track the environment with high temporal resolution^41^, to probe whether a central, modality-independent computation in higher-order cortex underlies temporal prediction.

Understanding the neural mechanisms of temporal anticipation is important as it constitutes a basic function that underpins many aspects of human perception and cognition. Building on our previous work^38,39^, we relate the foundational computation of event probability over time, as faithfully captured by our recently proposed model of temporal anticipation, to neural data.

## Results

We investigated neural activity during temporal anticipation with magnetoencephalography (MEG) while participants performed a Set-Go task in audition and vision (Fig. 2a). In the task, a Set cue was followed by a Go cue. The time between Set and Go, the *Go time*, was randomly drawn from truncated exponential or flipped exponential distributions, the *Go time distributions*, in separate blocks of trials (Fig. 2b). Participants were asked to press a button as fast as possible in response to the Go cue, generating RTs. In the task, fast reactions (short RT) represent strong event anticipation based on probability estimation, linking the experiment to many everyday tasks that require fast actions based on the prediction of future events.

**Fig. 2.**
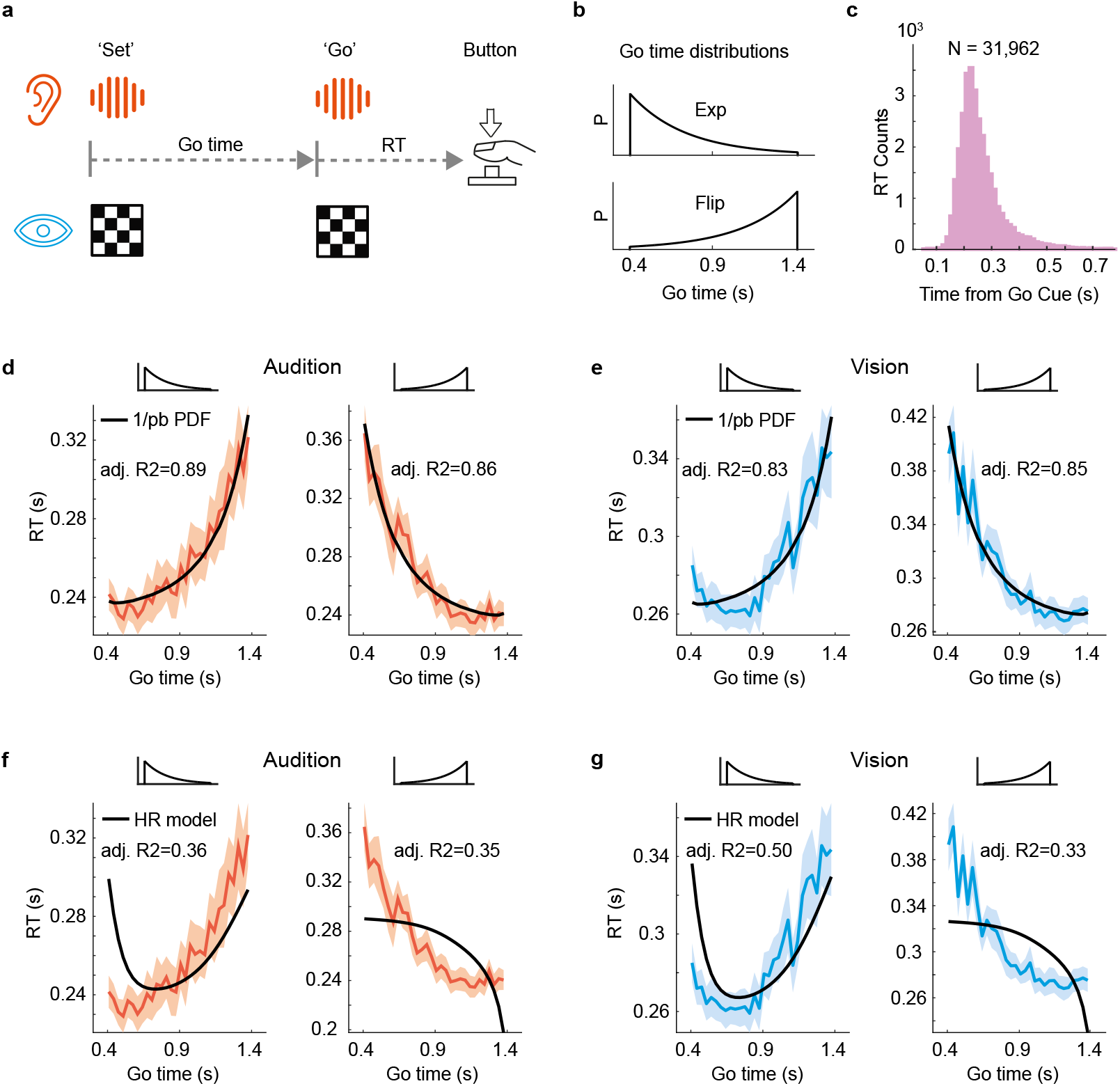
Participants estimate the probability density function of future events: temporal anticipation. **a**. Set-Go task schematic. In auditory and visual blocks of trials, participants were asked to respond as fast as possible to the Go cue, generating a reaction time (RT). In auditory blocks of trials, white noise bursts of 50 ms duration served as Set and Go cues; in visual blocks of trials, checker boards of 50 ms duration served as cues (Methods). **b**. In separate blocks of trials, the time between Set and Go (Go time) was drawn from a truncated exponential distribution (Exp) or from its left-right flipped version (Flip). **c**. Histogram of RT, pooled across the 23 participants and all experimental conditions. **d**. Reciprocal probabilistically blurred PDF fits mean auditory RT over Go time (Methods). **e**. Reciprocal probabilistically blurred PDF fits mean visual RT. **f**. Mirrored temporally blurred HR (Methods) does not capture auditory RT. g. Mirrored temporally blurred HR does not capture visual RT. **d-g** Shaded areas represent s.e.m. across participants.

23 participants performed the Set-Go task and generated 31,962 RTs (Fig. 2c). The RTs displayed typical features of a simple RT task^30^: the RT histogram showed a steep left flank and a short right tail, mean RT was short with small variance (0.261 s, SD = 0.085 s), 97.3 % of RTs fell inside of the interval RT = [0.1, 0.5] s. Mean RT was shorter in the auditory than in the visual conditions (−0.032 ± 0.031 s, mean ± SD, *P* = 0.0076, t = −2.8).

The Go time distributions modulated auditory and visual RTs in a systematic way: when Go cue probability was large, RT was short and vice versa (Fig. 2d and e). As Go cue probability decreased over time, RT increased (exponential conditions, Fig. 2d, left and 2e, left) and vice versa (flipped exponential conditions, Fig. 2d, right and 2e, right). The similarity of these RT dynamics across audition and vision suggests similar computations underlying the generation of RT in both modalities.

### Models of temporal anticipation based on probability estimation

For a long time it has been postulated that the critical variable underlying temporal anticipation, i.e. relating stochastic sensory input to RT output, is the HR of events^11-17,19,20,32,33^. Recent work has challenged this hypothesis in simple temporal anticipation tasks by demonstrating that humans estimate the event PDF but not the HR^38,39^. Here, both hypotheses were examined in order to verify that the better fit of the PDF-based model over the HR-based model to the behavioral data is replicated.

Importantly, the functional form of the model of anticipatory behavior has specific implications for underlying neural dynamics. The HR-based model accounts for two sources of uncertainty with specific hypotheses about their functional form. The first source is the uncertainty in time estimation which is hypothesized to linearly increase with elapsed time (“scalar property”)^42,43^. This hypothesis follows a simple intuition: the accuracy of time estimation linearly decreases with the length of the to-be-estimated time interval. The second source of uncertainty is the distribution of the event occurrence probability across time, the event PDF.

The brain’s estimate of the event PDF is hypothesized to be affected by the uncertainty in elapsed time estimation. In modeling, this is instantiated by a convolution of the objective event PDF with a Gaussian blurring kernel representing the uncertainty in elapsed time estimation, a computation termed *temporal blurring* (Fig. 3a, left).

**Fig. 3.**
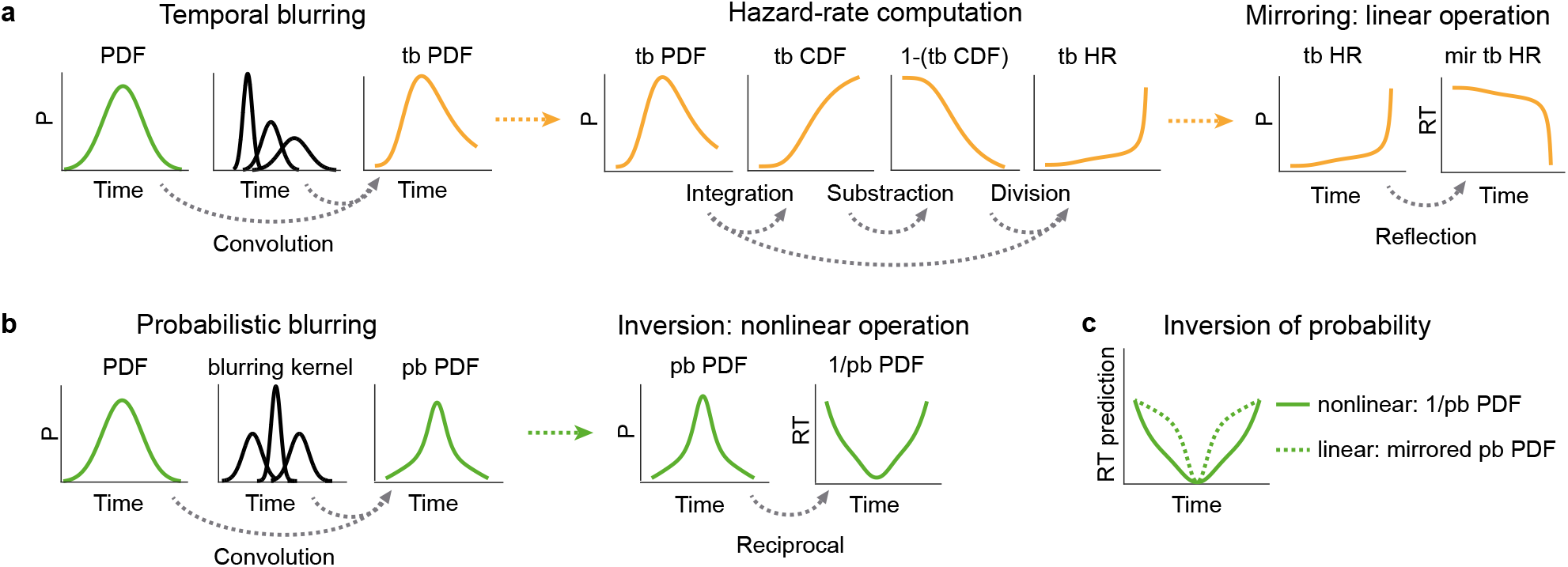
Computational hypotheses in temporal anticipation. **a**. Canonical hazard rate hypothesis. Left: temporal blurring (“tb”) poses that uncertainty in neural estimation of elapsed time increases linearly with elapsed time, affecting the estimate of event PDF (Methods). Middle: HR computation requires a sequence of three mathematical operations (Methods). Right: HR hypothesis predicts a linear, mirrored (“mir”) relationship between event probability over time and anticipation (Methods). **b**. PDF-based hypothesis. Left: Probabilistic blurring. Event PDF determines uncertainty in elapsed time estimation. In modeling, the objective event PDF is convolved with a Gaussian kernel whose SD is inversely related to the event PDF, resulting in a subjective estimate of the event PDF: the probabilistically blurred (“pb”) PDF (Methods). Right: *Reciprocal* probabilistically blurred PDF (“1/pb PDF”) predicts RT. **c**. Effect of reciprocal computation on RT prediction. 1/pb PDF implies a nonlinear “weighting” of event probability in temporal anticipation. For comparison, probabilistically blurred PDF was linearly inverted (“mirrored pb PDF”) to illustrate the effect of the presumed nonlinear reciprocal computation. For comparison, both 1/pb PDF and mirrored pb PDF were scaled by their respective ranges. Y axis in arbitrary units.

The HR-hypothesis implies that neural systems transform the temporally blurred event PDF to a more complex form, the Hazard Rate (HR). Mathematically (and computationally), this requires three distinct sequential steps: First the estimation of the PDF, *f*(*t*), then the estimation of its integral across time for computing the cumulative density function (CDF), 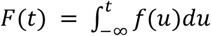 and, lastly, subtraction and division for estimating the HR as 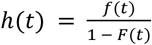 (Fig. 3a, middle).

Finally, the HR model hypothesizes an inverse relationship between the neural representation of HR and a motor response; i.e. when HR assumes large values, anticipation is strong, predicting short RT and vice versa. This relationship is expressed by a linear mirroring of the HR variable around a fixed value (Fig. 3a, right).

The PDF-based model has three main contrasting assumptions to the HR model. First, it postulates that the human brain represents the event probability across time in a computationally simpler form than the HR, i.e. by estimating the PDF itself. Second, in the PDF-based model, the uncertainty in elapsed time estimation does not increase monotonically with time but it is modulated by the event probability density: when event probability is large, temporal estimates are precise and vice versa. This concept is termed *probabilistic blurring*^*38,39*^ (Fig 3b, left, Methods). Third, the inverse relationship between probability and RT is assumed to be reciprocal^38,39^ (Fig. 3b, right).

### Computation of event PDF underlies temporal anticipation behavior

Both the HR-based variable (temporally blurred mirrored HR, Methods) and the PDF-based variable (probabilistically blurred reciprocal PDF, Methods) were computed and fit to RT curves. The PDF-based model captured the RT dynamics well in audition (Fig. 2d) and vision (Fig. 2e), and in both exponential and flipped exponential conditions, as supported by large values of adjusted R^2^. In contrast, the fits of the canonical HR-based model failed to capture the RT dynamics (Fig. 2f and 2g). Each HR-based fit resulted in a substantially smaller value of adjusted R^2^ compared to the PDF-based model. In the flipped exponential case the HR-based model even made the qualitatively wrong prediction of a convex RT shape where the RT curve was concave (Fig. 2f and g, right).

There was no evidence from behavior that humans compute the HR to predict the timing of future events. Instead, the analysis of the psychophysical data indicates that participants estimate a computationally simple variable, the event PDF, to predict the timing of future sensory cues, perfectly replicating previous findings^38,39^.

### Alpha and beta power decrease prior to an anticipated event

We sought to identify the neural signatures of anticipation in cortical population activity using MEG. We focused the analysis on the time span preceding the Go cue (Fig. 2a) since this is when a representation of the event PDF should be reflected in neural activity.

The MEG data were aligned to the Set and to the Go cues, respectively. In each alignment, the time-frequency decomposition was computed at each time point of each trial using a moving window (Hanning). We first investigated power changes prior to the Go cue relative to a pre-Set cue baseline (−0.4 to 0 s).

Before the Go cue, power decreased significantly in the alpha (7–12 Hz) and beta (15–35 Hz) frequency bands (Fig. 4a, P = 0.001, cluster test, Methods). Such differences in neural activity between pre-Set and pre-Go periods suggest activity related to Go cue anticipation, based on task demands.

**Fig. 4.**
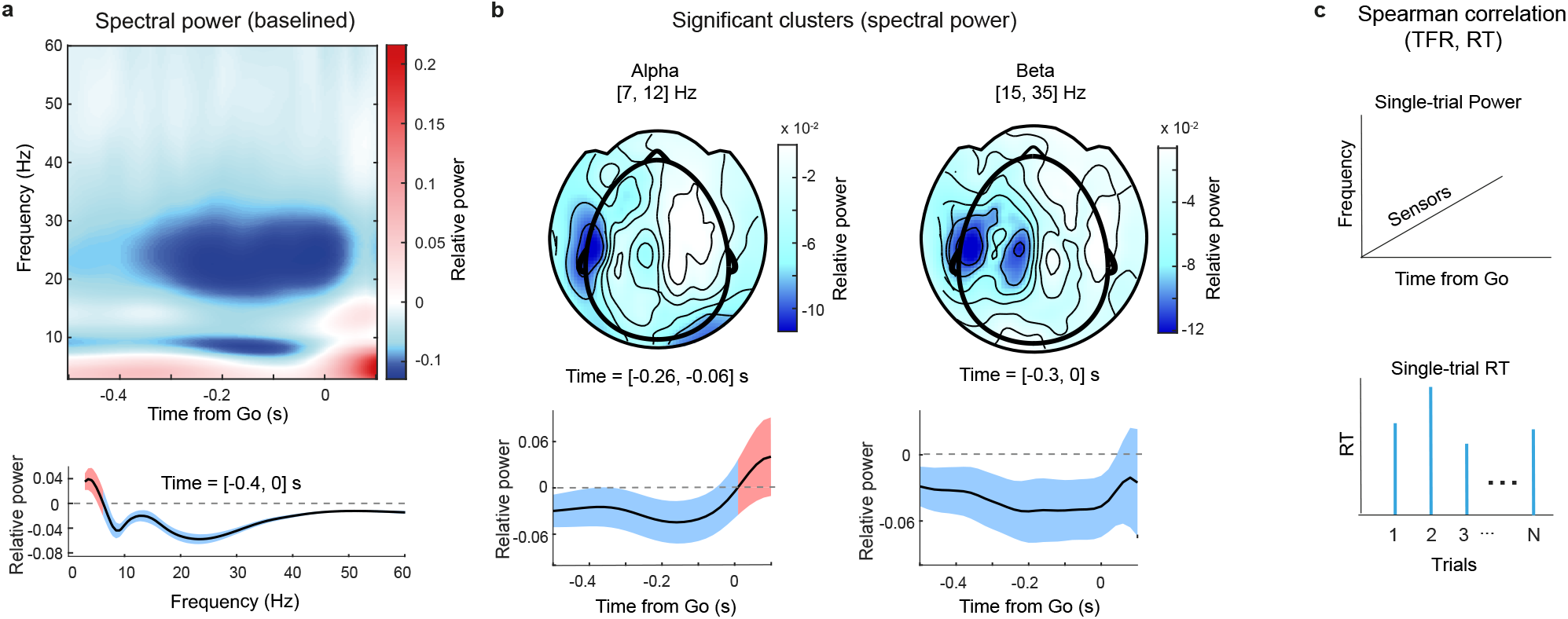
Alpha and beta band power decrease before anticipated events. **a**. Top: Spectral power averaged across participants, conditions (audition and vision, exponential and flipped exponential distributions) and MEG channels, baselined relative to a pre-Set cue time span (t = [−0.4, 0] s, Methods). Desynchronization in alpha (7–12 Hz) and beta (15–30 Hz) bands prior to Go cue. Bottom: Decrease in alpha and beta spectral power changes averaged across channels and time (t = [−0.4, 0] s), shaded area depicts SD across time. **b**. Top: Sensor array topography of significant desynchronization in alpha (left) and beta (right) power (cluster test, power against zero). Bottom: power change averaged across channels, shaded areas depict SD across channels. **c**. Correlation analysis schematic: within each experimental condition, single-trial Spearman correlation is computed between power in each time-frequency-sensor triplet (not baselined) and RT.

The relative decrease in alpha power was located in two areas of the sensor array (Fig. 4b, top left). One location comprised sensors over left motor areas, in line with sensorimotor alpha (mu) ^44^. This raises the question whether sensorimotor alpha plays a role in temporal prediction which we address later with a correlation analysis. The other location comprised right posterior sensors. In the case of *spatial attention*, a lateralization of posterior alpha power is commonly reported, i.e. power increases ipsilateral to the direction of attention (expected stimulus direction) and decreases contralaterally. This is often observed in visual^45^ and, in a similar fashion, in auditory tasks^46^. The present study, however, does not feature spatial lateralization as a component: stimulation in vision and audition was bilateral. Therefore, the rightward lateralization observed here does not indicate a spatial attention phenomenon.

Alpha power was consistently suppressed prior to the event, with the strongest desynchronization around 150 ms before the Go cue. Interestingly, alpha suppression was progressively diminished after that, reaching zero at the time of the Go cue (Fig. 4b, bottom left). These dynamics are consistent with an anticipation signal.

The relative decrease in beta power was located in sensors over left motor areas (Fig. 4b, top right). Beta power was also consistently suppressed before the Go signal, with the maximum suppression occurring during the last 200 ms an then returning towards zero. Again, this suggests an association to prediction (Fig. 4b, bottom right).

Finally, we observed an increase of low-frequency power (3–7 Hz, Fig. 4a) just after the Go cue occurrence, which reflects the signatures of the event-related field transient activity^47^ which also extended to higher frequencies.

### Sensor-level correlation analysis scheme

To investigate how the observed alpha and beta band neural dynamics relate to the temporal anticipation of the Go cue, we computed Spearman’s correlation between power in each MEG time-frequency-sensor triplet (Fig. 4c, top) and RT at the single-trial level (Fig. 4c, bottom, Methods). A cluster-based permutation test on Spearman’s rho identifies clusters in time-frequency-sensor space in which rho significantly differed from zero. Importantly, this cluster test was run across the two modalities (audition and vision) and the two Go time distribution conditions (exponential and flipped-exponential) in order to reveal correlation clusters shared by all experimental conditions.

### Correlation between spectral power and RT indicates a neural representation of event PDF

We employed a correlation-based analysis strategy that identifies neural activity *prior* to the Go cue that is correlated with RT. Note that RT demarcates the time point of the button press *after* the Go cue. Thus a significant correlation indicates that neural activity *before* the Go cue is *predictive* of RT *after* the cue. Since RT is driven by event probability, as demonstrated by the convincing fits of the PDF-based RT model, the correlation between pre-Go-cue spectral power and RT can be interpreted as a neural representation of the event PDF.

### Sensor-level alpha and beta power represent event PDF

The correlation analysis between spectral power and RT identified a negative correlation cluster that comprised both alpha and low beta bands, namely the range between 7 to 22 Hz (Fig. 5a, top, P = 0.001). Notably, the dynamics of Spearman’s rho over frequency (Fig. 5a, bottom), with two distinct peaks (troughs) in the alpha and low beta ranges, support a within-cluster separation into these two bands. Accordingly the cluster was split into an alpha and a beta bands, with frequency ranges 7 to 12 Hz and 15 to 22 Hz respectively. In each of these two frequency bands, only the significant sensors within the corresponding frequency band were selected.

**Fig. 5.**
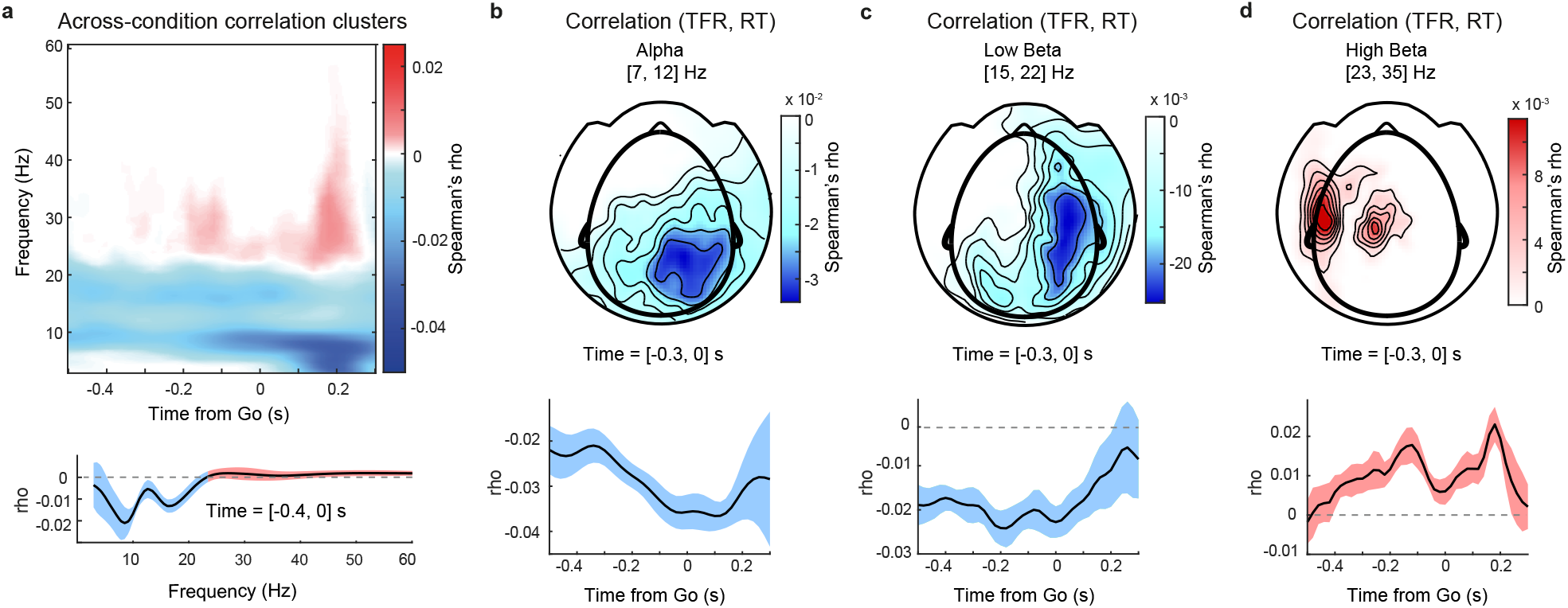
Sensor-level alpha and beta power predict reaction time before anticipated events. **a**. Top: Negative correlation cluster comprises alpha and low beta frequency bands (P = 0.001), positive correlation cluster in high beta band (P = 0.087) (across-condition cluster tests, Methods). Bottom: Mean Spearman’s rho across time and sensors, shaded area depicts SD across time. **b**. Top: Sensor-level topography of alpha band correlation. Bottom: Mean rho and SD across sensors. Before Go cue, rho decreases until right before Go cue **c**. Sensor-level topography of low beta band correlation. Bottom: Mean rho and SD across sensors. Rho increases right after Go cue. **d**. Sensor-level topography of high beta band correlation cluster. Bottom: Mean rho and SD across cluster sensors. Rho increases until right before Go cue, then decreases reaching a local minimum at time of Go cue.

Additionally, a positive correlation cluster was observed in the high beta (23–35 Hz) band (Fig. 5d, P = 0.087, cluster test).

Topographically, the negative alpha band correlation was right lateralized over right posterior sensors (Fig 5b, top). No corresponding negative correlation was found over the left sensorimotor areas, which shows that sensorimotor mu rhythm was not modulated by anticipation, as hypothesized above (Fig. 4b, top left). Within the alpha frequency channels, in the time span before the Go cue, Spearman’s rho decreased over time from −0.3 s to 0 s and reached its largest negative values immediately before the Go cue (Fig. 5b, bottom). This indicates that alpha power was most predictive of RT at the time point of Go cue occurrence, suggesting a functional role of alpha oscillations in temporal anticipation.

The negative correlation in the low beta band was also located over right posterior sensors and extended towards right frontal sensors (Fig. 5c, top). Between −0.4 and 0 s relative to the Go cue, Spearman’s rho remained consistently negative. Immediately after the Go cue, rho increased towards zero (Fig. 5c, bottom), similar to the correlation in the alpha band. This dynamic entails that low beta power is most predictive of RT right until Go-cue occurrence, hinting towards a functional role of low beta oscillations in temporal anticipation.

In the high beta band, the positive correlation cluster was left lateralized and was located over left sensorimotor areas (Fig. 5d, top). Spearman’s rho, averaged across the cluster’s sensors, displayed an interesting dynamic: Rho increased from −0.4 to −0.1 s relative to the Go cue, peaked at around −0.1 s, and thereafter rapidly decreased, reaching a minimum at the time-point of Go cue occurrence (Fig. 5d, bottom). At ∼0.2 s after the Go cue, rho reached its peak and rapidly decreased towards zero thereafter. In light of the distribution of RTs (Fig. 2c), which has a grand mean of RT = 0.261 s, the correlation coefficient peaks ∼60 ms before the button press, suggesting that the modulation of high beta power is related to motor execution. This interpretation is in line with the common finding of a beta band desynchronization during preparation and execution of movements^48^.

Taken together, the sensor-level analyses identified three frequency bands-of-interest in which spectral power prior to the Go cue was correlated with RT. Importantly, two of the neural signals seem linked to prediction: alpha band correlation decreased and high beta band correlation increased *until just before* the Go cue, suggesting a functional relationship between spectral power and temporal anticipation. Low beta band correlation also displayed an interpretable dynamic: its power was most predictive of RT before Go cue occurrence. The fact that spectral power in the three frequency bands *before* the Go cue is correlated with RT, i.e. the button press *after* the event, indicates that a neural representation of the event PDF underlies the anticipation of sensory cues. In order to better understand the functional role of these neural dynamics, we next aimed to reveal the cortical localization of the three correlation signals.

### Source-level correlation analysis scheme

Within each frequency band-of-interest (7–12 Hz, 15–22 Hz, 23–30 Hz), identified in the above sensor-level analysis, spectral power was projected to brain source-space using a spatial filter (DICS beamforming^49^, Methods). This source-space for each participant comprised the individual’s cortical mantle representation, as extracted from a structural MR scan. This source space was fused with a whole-cortex parcellation atlas (AAL atlas^50^, Methods) in order to relate findings to known anatomical areas and assist the interpretation of results. Within each brain source, power in each frequency band-of-interest was averaged across its frequencies. Spearman correlation was then computed between single-trial average power and single-trial RT in each source separately. Finally, a cluster test across all experimental conditions (audition and vision, exponential and flipped exponential Go time distributions) identified clusters of cortical sources in which Spearman’s rho significantly differed from zero. This across-condition approach identifies correlation signals common to all experimental conditions. To investigate correlation dynamics across time, this analysis was run within each of eight time-windows relative to the Go cue, centered at t = −0.35 to t = 0.35 seconds in steps of 100 ms (Methods).

### Alpha band power in right IPL and pMTG region predicts RT to anticipated events

The source-level correlation between alpha (7–12 Hz) power and RT was right lateralized (Suppl. Fig. 1), covering posterior parietal and posterior temporal cortices (Fig. 6a top row). Spearman’s rho decreased prior to the Go cue and reached the largest negative values around 50 ms before the cue (Fig. 6a top row, Fig 6b), confirming the dynamics observed in sensor-level analysis (Fig. 5b, bottom). After 0.15 s post cue, there were no significant clusters.

**Fig. 6.**
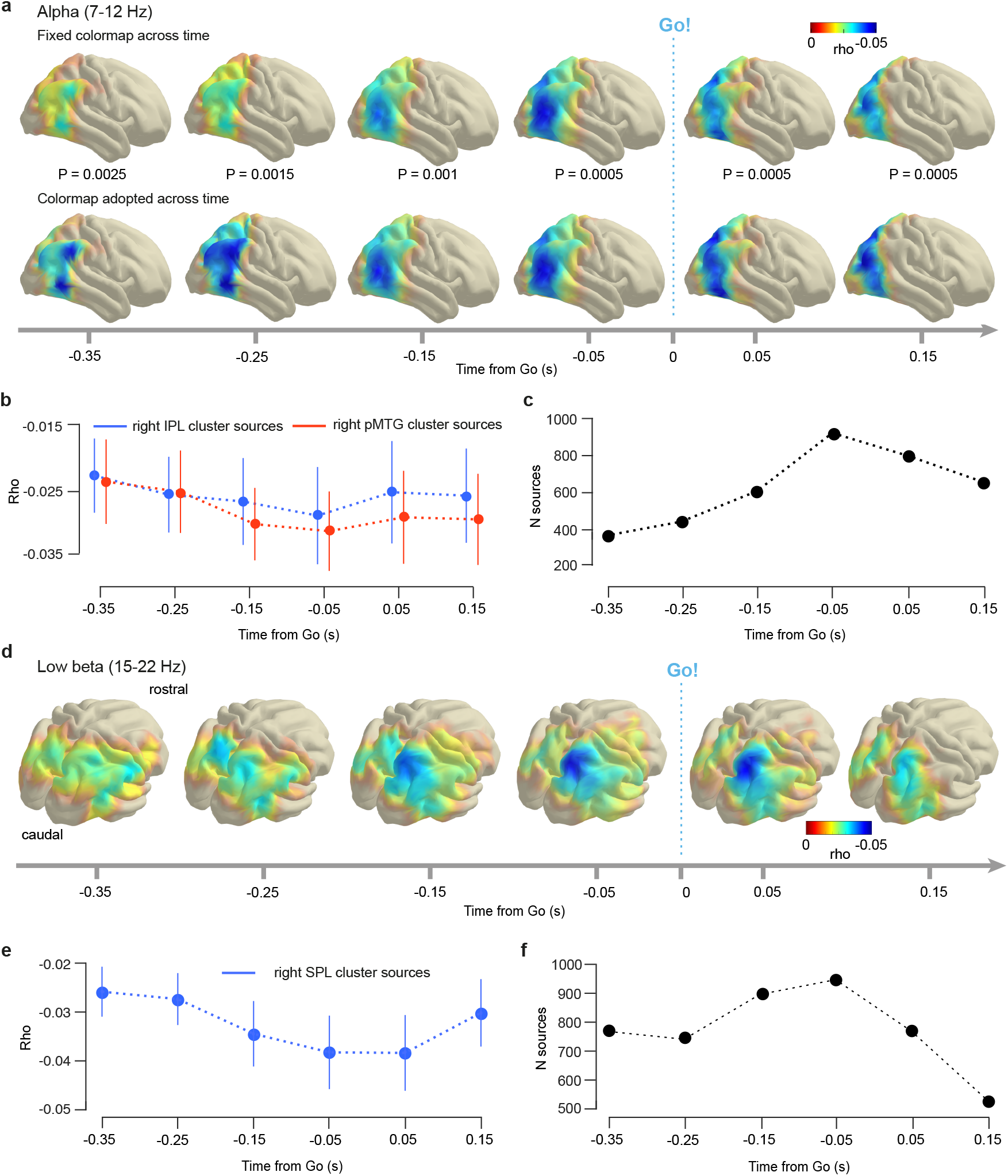
Alpha band power in right parieto-temporal areas and low beta band power in right superior parietal lobule predict reaction time to anticipated events. **a**. Top row: Cortical topography of across-condition Spearman correlation (mean alpha (7–12 Hz) power, RT). Negative correlation reaches maximum in posterior middle temporal gyrus (pMTG) region immediately before the Go cue. Colormap fixed across time to highlight across-time correlation dynamics. Bottom row: Re-plotting of correlation shown in top row with an individual colormap per plot (Suppl. Fig. 2) highlights correlation extent for each time span. Negative correlation at −0.35 s in supramarginal gyrus and pMTG spreads out to cover entire inferior parietal lobule (IPL) at −0.25 s and extends to pMTG region from −0.15 s on. **b**. Mean Spearman’s rho averaged across sources from two regions-of-interest (blue: right IPL, red: right pMTG). Averaging first done within-participant and across sources, then across participants. Error bars are standard error of the mean. **c**. Number of sources over time. **d**. Cortical topography of across-condition Spearman correlation (mean low beta (15–22 Hz) power, RT). Negative correlation is maximal in right superior parietal lobule (SPL) before the Go cue. All colormaps fixed across time to highlight across-time correlation dynamics. **e**. Mean Spearman’s rho averaged across sources from right SPL. Averaging first done within-participant and across sources, then across participants. Error bars are standard error of the mean. **f**. Number of sources over time. P values uncorrected.

To demonstrate more precisely the anatomical distribution of the correlation signal over time, we re-plotted the same analysis using an individual colormap for each time window (Fig. 6a bottom row, Suppl. Fig. 2). This revealed that at −0.35 s before the Go cue, the correlation signal has prominent local maxima in right supramarginal gyrus (SMG) and posterior middle temporal gyrus (pMTG). Beyond this early time window, the signal quickly spread out across the entire right inferior parietal lobule (IPL) at ∼-0.25 s and extended to the posterior MTG region at −0.15 s. After the Go cue, the correlation decreased in cortical extent (Fig. 6c). This source-level analysis thus identified two cortical regions, one including the right IPL, the other the right pMTG, in which the dynamics of alpha band oscillations followed an approximation of the event PDF before the occurrence of an anticipated event. These findings suggest a supra-modal, functional role of the right IPL and right pMTG area in temporal event anticipation.

### Low beta band power in right SPL predicts RT to anticipated events

The source-level correlation between low beta power (15–22 Hz) and RT was right lateralized in six time windows spanning −0.35 to 0.15 s relative to the Go cue (Fig. 6d). In this time range, significant negative correlation (all P values = 0.0005, uncorrected, Methods) covered parts of right PPC but not occipital cortex. Notably, the correlation signal was strongest in right superior parietal lobule (SPL) at −0.05 s, i.e. just before Go cue occurrence (Fig. 6e and f). There was no significant correlation after 0.15 s. The analysis identified right SPL as the cortical site where low beta power is most predictive of RT before the anticipated cue. This relationship between low beta oscillations and RT indicates a representation of the event PDF in cortical dynamics, suggesting a supra-modal, functional role of right SPL in temporal anticipation.

### High beta band power in left sensorimotor cortex predicts RT to anticipated events

The positive correlation in the high beta band (23–30 Hz) were localized in left sensorimotor cortex (Fig. 7a), in agreement with the right index finger button press. The correlation signal increased in magnitude over time, peaked at 0.05 s post cue and decreased thereafter (Fig. 7b). The cluster extent, expressed as number of brain sources, displayed a similar dynamic (Fig. 7c), it increased prior to the Go cue and decreased quickly thereafter.

**Fig. 7.**
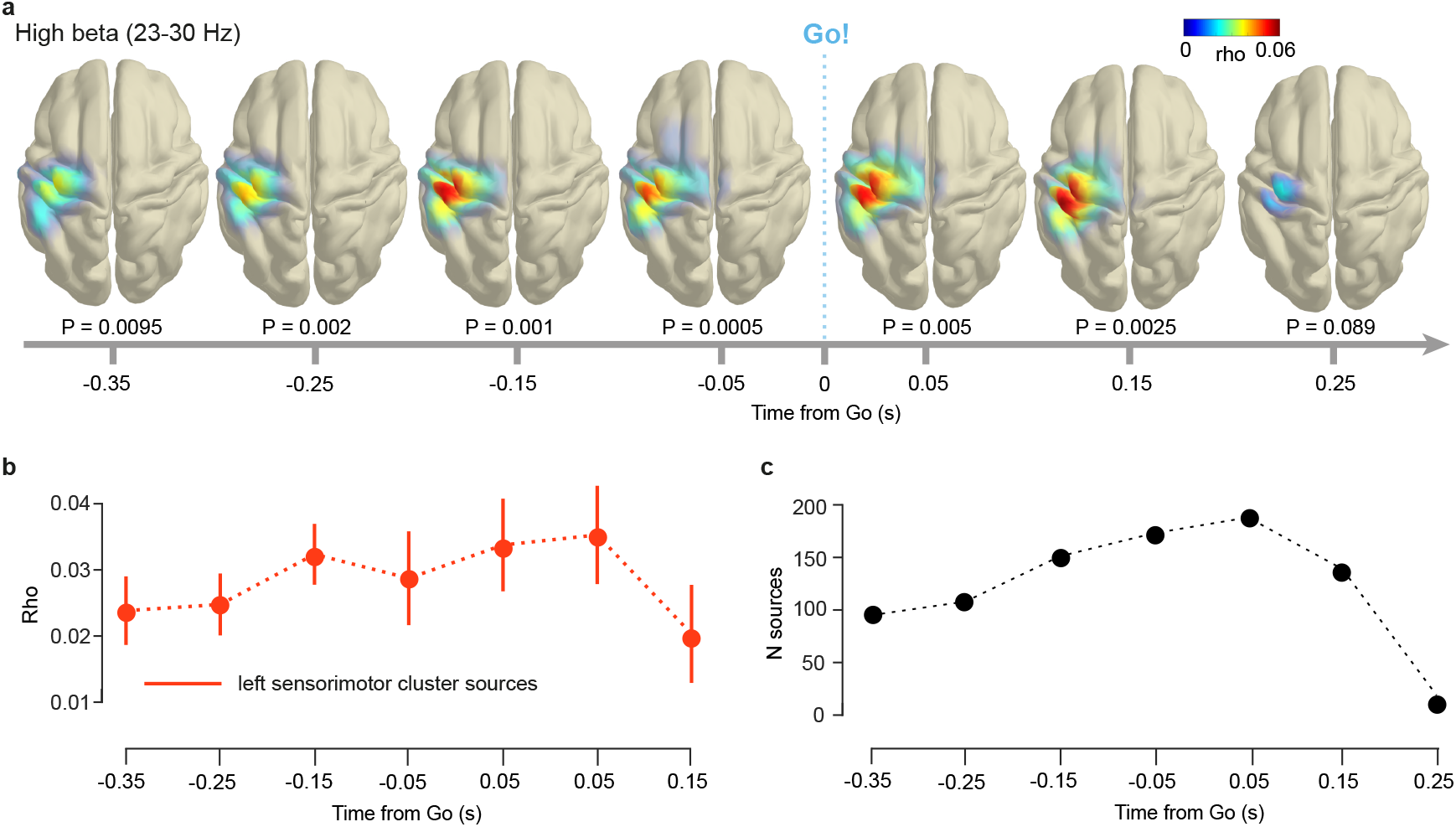
High beta band power in left sensorimotor areas predicts reaction time to anticipated events. **a**. Cortical topography of across-condition Spearman correlation (mean high beta (23–30 Hz) power, RT. Positive correlation in left sensorimotor cortex prior to Go cue. All colormaps fixed across time to highlight across-time correlation dynamics. **b**. Spearman’s rho averaged across sources from left sensorimotor cortex. Averaging first done within-participant and across sources, then across participants. Error bars depict standard error of the mean. **c**. Number of sources over time. P values uncorrected.

The positive sign of the correlation signal is in line with a hypothesized disinhibitory role of beta band desynchronization^48^: small power values predicted short RTs and vice versa. This interpretation is supported by the fact that the correlation signal increased before participants pressed the response button. Interestingly, the positive correlation between RT and beta power occurred hundreds of milliseconds before the anticipated sensory event (Go cue), demonstrating that motor preparation itself follows an approximation of the event PDF.

The correlation analysis between spectral power and RT reveals that, before anticipated auditory and visual sensory events, alpha, low beta and high beta spectral power dynamics represent an approximation of the event PDF. The cortical location of these representations in right IPL and pMTG (alpha), right SPL (low beta), and left sensorimotor cortex (high beta) invite a functional interpretation, suggesting temporal attention (alpha), motor preparation (low beta), and motor disinhibition (high beta), as will be developed in the Discussion. Taken together, these three main results link the specific computation of event PDF to right-lateralized cortical dynamics and demonstrate a supra-modal role of cortical areas SPL, IPL, pMTG, as well as left sensorimotor cortex in the prediction of future events.

### A neural representation of the probability density function of sensory events

In an additional time-frequency decomposition analysis, we aimed to identify a direct representation of the event PDF in neural dynamics. Based on the convincing fits to group-level RT (Fig. 2d and 2e), we fit the PDF-based variable (probabilistically blurred reciprocal PDF) to single-trial RT (Methods). This PDF-based model of RT was then Spearman-correlated with single-trial source-level power, averaged within each of the three frequency bands-of-interest (7–12 Hz, 15–22 Hz, 23–30 Hz). A cluster-based permutation test identified the sources in which Spearman’s rho significantly differed from zero (Methods). This correlation analysis was performed within four time windows relative to the Go cue (−0.4 to 0 s, window size 0.1 s).

The source-level correlation between power in the three frequency bands-of-interest and the PDF-based model of single-trial RT identified significant clusters. Remarkably, the analysis replicated the cortical topography of the correlation analysis between spectral power and RT reported above. In the alpha (7–12 Hz) band case, the analysis identified sources in right IPL and the pMTG area (Suppl. Fig. 3a). In the low beta (15–22 Hz) case, the correlation signal focused in the right SPL region (Suppl. Fig. 3b). In the high beta (23–30 Hz) case, the correlation signal covered left sensorimotor areas (Suppl. Fig. 3c). This analysis further demonstrates that neural dynamics in alpha and beta frequency bands represent the event probability distribution over time prior to anticipated sensory events in higher-order cortex and in sensorimotor cortex.

### Sensor-level ERF analysis scheme

The analysis of spectral power did not identify prediction signals in auditory or visual sensory cortex. Given the evidence for the involvement of early sensory cortex in prediction^10,51,52^, we next investigated potential effects of event probability density on neural activity time-locked to the Go cue, i.e. on the event-related fields (ERFs). The Go stimulus resulted in a prominent P1 response in both auditory (Fig. 8a top) and visual (Fig. 8a bottom) conditions, originating in primary sensory areas (topography plots, Fig. 8a). At the level of averaged data and based on visual inspection, the time course of early neural activity (< 0.15 s) was similar across the exponential and flipped-exponential probability conditions in both vision and audition. We next aimed to quantify potential neural signatures of event probability density in time-locked data *before* the Go cue and in early and late ERF components *after* the Go cue.

**Fig. 8.**
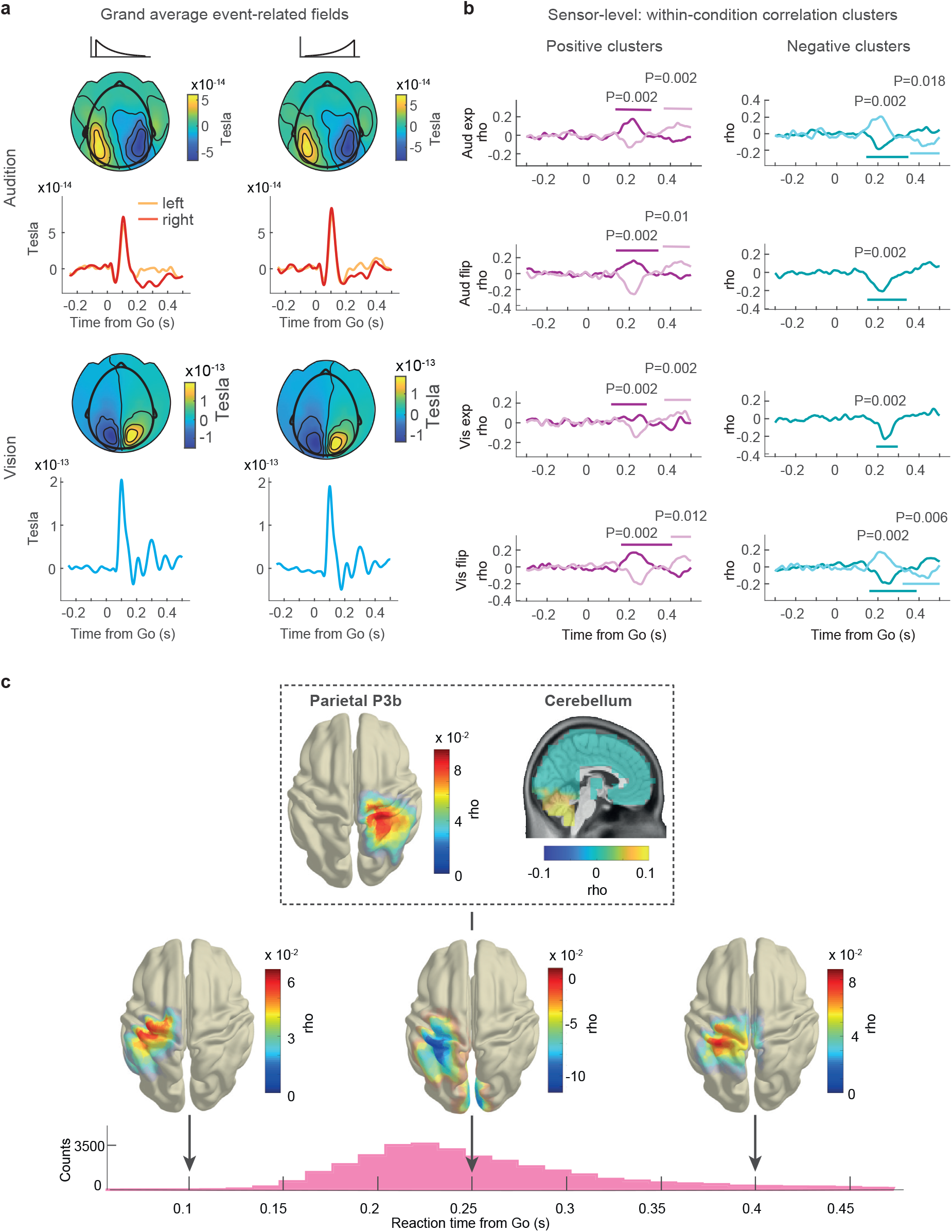
Parietal, cerebellar, and sensorimotor event-related fields correlate with reaction times to anticipated events. **a**. Grand-average event-related fields plotted through participant-level selection of ERF sensors over auditory and visual areas. Top: audition, bottom: vision. **b**. Early and late positive (left) and negative (right) sensor-level correlation clusters (ERF, RT). Mean Spearman’s rho averaged across clusters’ sensors over time. Colored lines demarcate cluster time spans, P-values uncorrected. **c**. Topography of across-condition source-level Spearman correlation clusters (ERF, RT). At 0.25 s post-Go cue, positive correlation clusters in right parietal cortex (P3b) and cerebellum (both top) and a negative cluster in left sensorimotor cortex (bottom). At 0.1 s and 0.4 s, positive correlation clusters in left sensorimotor cortex. Bottom: RT histogram from Fig. 2c for orientation.

We used a correlation-based approach to relate neural activity, time-locked to the Go cue, to RT. At the single-participant level, the MEG data and the RTs were aggregated within adjacent pairs of Go times (called *frames* from this point on) for a mild smoothing. For each sensor-by-time-point duplet, Spearman’s rho was computed between the 30 frames of MEG data and the 30 averaged RTs, resulting in a correlation matrix (sensors by time points). On all participants’ rho values, a cluster-based permutation test was run within each experimental condition to identify sensor-by-time clusters in which rho differs from zero (Methods). This analysis pipeline was run using a pre-Set baseline (−0.5 to 0 s) to investigate the time span *before* the Go cue, and, using a pre-Go baseline (−0.5 to 0 s), to investigate the time span after the *Go-cue*.

### Sensor-level event-related fields correlate with RTs

The analysis did not identify a significant correlation cluster in the time span *before* the Go cue. No correlation was observed in early sensory ERF components (<75 ms) in visual or auditory areas, indicating that either these sensory responses were not modulated by probability density or that our design was not sensitive enough to isolate these small effects which may require a dedicated design that allows for averaging across several hundreds of trials per frame^53^.

*After* the Go cue, two *positive* and two *negative* correlation clusters were identified. An early positive cluster (∼0.25 s post-Go) and a late positive cluster (∼0.4 s post-Go) occurred in all four experimental conditions (Fig. 8b, left). An early negativ*e* correlation cluster was identified (∼0.25 s post-Go) in all four conditions and a late negative cluster (∼400 ms post-Go) in the auditory exponential and the visual flipped exponential conditions (Fig. 8b, right). In order to better interpret the correlation, we selected two time points of interest (0.25 s and 0.4 s), that were shared by the positive and negative correlation clusters, for a source-level analysis.

### Source-level ERF analysis scheme

In a next analysis, we aimed to obtain a more precise spatial estimate of the correlations between ERFs and RT to better understand the functional role of time-locked neural activity in temporal anticipation. We employed a source-level correlation-based analysis. The per-frame MEG data (see above) were projected to source-space (LCMV beamforming, Methods) and Spearman’s rho was computed between ERF and RT for each source and time point. Rho was averaged across time within windows-of-interest that were chosen based on the sensor-space clusters (30 ms windows centered around 0.25 and 0.4 s post Go) and one additional 30 ms window centered at 0.1 s post Go to probe for early correlations before the button response. We carried out *across-condition* cluster-based permutation tests to reveal source-level correlation clusters shared by audition and vision.

### Parietal P300, motor, and cerebellar ERFs are sensitive to event probability density

We observed a significant positive cluster over right parietal cortex at t = 0.25 s post-Go (Fig. 8c, top left, P = 0.012). This cluster’s largest rho values covered a right parietal area that included right postcentral gyrus, anterior-lateral SPL, and intraparietal sulcus. The cluster’s temporal and anatomical locations are in agreement with the ERP component P300, more specifically, with the P3b which originates from parietal areas^53,54^. The positive correlation with RT implies that the P3b’s amplitude negatively covaries with probability density: P3b amplitude is small where probability is high and RT is short, and vice versa. This is in agreement with the P3b’s sensitivity to probability^55,56^, more specifically, with work that promotes a *negative correlation* between P300 amplitude and probability^57^, and our results extend these findings to probability *over time*.

At 0.25 s post Go, we identified a second positive correlation cluster consisting of sources in the cerebellum (Fig. 8c, top right, *P* = 0.005). The positive correlation expresses that when cerebellar activity is small, RT is short and vice versa. This finding is in agreement with the known functional role of the cerebellum in (finger^58^) movement control^59^. Alternatively it may reflect activity related to the representation of time, an activity that is also linked to the cerebellum^60^. Both interpretations seem plausible, but a targeted experiment is needed to assess more conclusively the function of the cerebellum in temporal anticipation.

At 0.1, 0.25, and 0.4 s post-Go cue, significant correlation clusters emerged over left sensorimotor cortex (Fig. 8c bottom, P = [0.028, 0.002, 0.012] uncorrected). Notably, the cluster at 0.1 s occured *before* the button was pressed (see RT histogram in Fig. 8c, bottom).

The positive correlation (small ERF amplitude, short RT and vice versa) likely corresponds to preparatory motor activity reflecting a state-of-readiness. The negative correlation cluster at 0.25 s post-Go was close to the grand mean of RT (0.261 s) and corresponds to the execution of the button press. This interpretation is supported by the negative sign of the correlation: the larger the ERF amplitude, the shorter RT and vice versa. The positive correlation cluster at 0.4 s post Go likely reflects a late component of the motor ERF: at 400 ms post-Go, a small amplitude corresponds to a short RT and vice versa. These clusters over left sensorimotor cortex were to be expected and provide a sanity check of the correlation analysis pipeline.

### Event probability density represented in parietal P300, motor, and cerebellar ERFs

In a final analysis, we aimed to identify a direct representation of the event PDF in neural activity time-locked to anticipated events. Using the source-level ERF analysis scheme reported above, we performed a source-level correlation between ERFs and fits of the PDF-based model to RT (Methods). This analysis confirmed the key results of the correlation analysis with RT: At 0.25s after the Go cue, parietal P3b and cerebellar ERFs and motor ERFs at 0.25s and 0.4s were correlated with the event PDF (Suppl. Fig. 4). This analysis revealed a representation of the probability distribution of sensory events in time-locked neural activity after an anticipated event.

In sum, the analysis of time-locked data identified significant correlations between ERFs and RT, as well as between ERFs and the PDF-based model of RT. The clusters over left sensorimotor cortex illustrate how effector motor activity relates to event probability density. The positive correlation clusters at 0.25 s post-Go demonstrate sensitivity of the parietal P3b and a cerebellar ERF to event probability density in temporal anticipation.

## Discussion

We investigated the neural correlates of temporal event anticipation. Reaction times to auditory and visual cues, distributed in time, were fit by a model based on the reciprocal event PDF. The canonical hazard-rate-based model failed to capture the RT data.

We report three main results. First, alpha power in the right IPL and pMTG area, low beta band power in the right SPL area, and high beta band power in left sensorimotor cortex represent the event PDF. These three anticipation signals occurred before the sensory event and predicted RT. Second, time-locked activity in right parietal cortex, after the anticipated event had occurred, was correlated with the event PDF, hinting at a functional role of ERF component P300 in temporal expectancy. Third, these results in time-frequency and time-locked neural data were observed in audition and vision, consistent with a supra-modal role of the three cortical areas in the representation of event probability density and in the generation of fast responses to predicted sensory events.

### The right PPC and pMTG in temporal anticipation

The integration of abstract information in sensorimotor transformation is a prominent PPC function. In action planning, this transformation incorporates information about *where* an object is in space and facilitates motor planning in e.g. reaching and grasping^27^. We investigated temporal anticipation, and the crucial information is *when* a future event will likely occur. Modulations of both low beta power in the SPL area and alpha power in IPL and pMTG represented event probability density in anticipation of the timing of future events. Interestingly, the right pMTG, a cortical area commonly associated with language processing^61,62^, exhibits a strong functional connectivity to the IPL (supramarginal and angular gyri) ^61^. The two signals carry the information needed to shape the moment-to-moment readiness-to-respond leading to the observed probability-sensitive reaction times. It is therefore a reasonable assumption that these signals affect the generation of a motor plan which is then executed in left motor cortex, resulting in a probability-modulated right index finger button press.

With respect to the functional role of the PPC in temporal anticipation, two major points require discussion: First, we address the strong rightward lateralization observed in the two prediction signals, then we consider potentially different functional roles of the two signals. A fundamental process demanded by our experimental task is the estimation of elapsed time relative to the Set cue that allows for orienting in time in anticipation of the Go cue. It is argued that the computational processes underlying such time estimation are directly linked to neural population dynamics^18^. At the (coarse) level of the cortical lobes, the representation of time between sensory events is often linked to parietal activity. In macaques, the posterior parietal cortex has been implicated in the representation of time^63^. In humans, activity in the right posterior inferior parietal lobe is related to event timing^64,65^, whereas the left parietal lobe does not seem to play such a critical role for processing of stimulus timing^66,67^. Recent work proposes the right supramarginal gyrus as the locus of the subjective experience of time^68^.

These close connections between basic timing aspects and right parietal areas link the right lateralization observed in our data to the processing of event probability over time in temporal anticipation.

One obvious question is how the observed lateralization to right parietal areas relates to the generation of a left lateralized motor command, as required by our task. The functional connectivity of SPL and IPL with other cortical areas is vast (see^69-71^ for a comprehensive overview). The PPC exhibits connectivity between SPL/IPL and ipsilateral premotor areas but also between corresponding SPL/IPL contralateral areas^69^. Specifically, there is a marked structural connectivity between right SPL via the corpus callosum to the contralateral PPC^70^. The functional connections of many parietal regions are stronger in the right than in the left hemisphere^69^, which is thought to be consistent with the important role of the right hemisphere in spatial processing. Likewise, our case of temporal processing suggests a right hemispheric prioritization, too. Indeed, PPC hosts a bimanual representation of the limbs^28^ and PPC neurons code for movements of the contralateral limb^29,72^. In general, the PPC’s connectivity seems appropriate to support time-critical motor commands for the right hand fingers driven by right PPC sensory-to-motor transformations.

We observed two different prediction signals, one with a focus on the right SPL, the other with a focus on the right IPL and pMTG. However, our experimental design does not allow for a conclusive assessment of the functional roles of each of these signals. Still, the anatomical segregation as well as the segregation into low beta and alpha band signals invites different functional interpretations with respect to attentional phenomena.

Temporal anticipation is closely linked the deployment of attention over time^12^. The right parietal lobe is associated with attention to brief visual stimuli^73^, auditory selective attention^74^, and temporal visual attention in general^66,75^. The right SPL is part of the dorsal ventroparietal attention network that is involved in the selection of stimuli in visual search and detection and hosts top-down signals in visual expectancy^76^. But this attention network is largely bilateral and not known to be right lateralized^76^. In light of the strong right lateralization observed in the current data, the signal in right SPL may be more related to motor preparation based on integration of temporal-probabilistic stimulus information.

In contrast, the right IPL is part of the ventral frontoparietal attention network that is strongly right lateralized and is functionally associated with the detection of behaviorally relevant, salient stimuli in visuospatial attention^76^. Such attention functions are closely associated with alpha band power modulation discussed in the next section. The above interpretations link the low beta band signal in the right SPL area more to motor functions and the alpha band signal in the right IPL/pMTG area more to attention in time. However, this interpretation remains speculative and dedicated experiments are needed for a more conclusive assessment.

### Alpha band spectral power in temporal anticipation

Prior to the anticipated Go cue, alpha spectral power was negatively correlated with Go cue probability density. Source analysis revealed the right IPL and posterior MTG areas as the main loci of this co-modulation, suggesting that the event PDF may be computed, or at least represented, in right parietal areas. This assumption is in line with the parietal lobe’s role in sensorimotor integration^77^ and action planning^27^. Still, it raises the question about the functional role of alpha oscillations in parietal cortex.

In earlier cortical stages, in visual discrimination experiments, an inverse relationship between alpha spectral power and perceptual performance is commonly reported^78,79^. This phenomenon may be mediated by alpha oscillations driving the excitability of cortical tissue where increased alpha power is associated with low excitability (inhibition) and decreased alpha power is associated with high excitability^80^. Here we demonstrate a negative correlation between parietal alpha power and RT, i.e. large alpha power values are associated with short RTs and vice versa. It is not readily apparent how this relationship can be reconciled with the functional hypothesis of alpha oscillations as an inhibition signal. Instead, the negative correlation points to the event-related synchronization (ERS) hypothesis, which predicts alpha synchronization when the execution of a response is withheld over cortical areas that are under top-down control^80,81^. According to this hypothesis, a functional interpretation of our findings would be that event probability density drives the state of excitability; e.g. when event probability density is high, alpha power increases, cortical excitability is low, and the preparation of a motor command is withheld, which may lead to a rapid response (short RT) once the stimulus is registered. In short, higher predictability may lead to stronger inhibition of pre-event neural processes, potentially resulting in better preparation of the perceptual-motor network to respond to an imminent event.

The negative correlation between RT and alpha band activity further suggests a potential top-down control mechanism, that in our case may be sensitive to event probability density. This interpretation is supported by recent work proposing frontal top-down control of right parietal cortex in the context of the frontoparietal attention network^82^.

In spatial attention, anticipatory alpha desynchronization is lateralized in (occipito-)parietal areas as reported in the case of unilateral visual stimulation^83,84^. In our experiment, we stimulated bilaterally in audition and vision, but alpha desynchronization was right lateralized where one would have expected a bilateral effect in spatial attention tasks. This points to a potential processing difference between spatial and temporal information.

In contrast to spatial attention tasks, our experimental task demands the estimation of elapsed time. In our task, the estimation of time is subjective, i.e. time is associated with uncertainty^11,39,43^. Recent work proposes the right SMG as a locus for the *subjective experience of time*^68^. In our analysis, the right SMG was part of the negative correlation clusters identified in the alpha band, suggesting a role of rSMG in time estimation in temporal anticipation. In light of the right parietal lobe’s association with attention, e.g. to brief visual stimuli^73^, auditory selective attention^74^, and temporal visual attention in general^66,75^, the co-modulation between alpha power and probability-driven RT may be interpreted as a *temporal* attention phenomenon.

While many studies link activity in the alpha band to sensory cortices, the current experiment relates it to higher-order cortex. We show a supra-modal representation of event probability density, located in right IPL and posterior MTG areas that may be tightly linked to time estimation, inhibition, and temporal attention.

### The parietal P300 in temporal anticipation

Our analysis identified a modulation of the parietal P300 (P3b) ERF component by event probability density. The fact that the P300 occurred *after* the button response raises the question of the component’s functional role in temporal anticipation. In more complex decision-making tasks involving choice between options, the P300 occurs *before* the choice is instantiated and is argued to (partly) represent decision processes^85^, such as evidence accumulation^86^. Our simple task only requires a decision regarding the question: “Shall I press the button now?” which is solely contingent on the occurrence of the Go cue. We therefore favor other functional interpretations, such as that the P300 may represent processes involved in the updating of working memory^87-89^ or information updating in general. In this regard, the concept of context updating^90^ proposes an attention-driven comparison between current sensory input and the representation of past sensory events. Our analysis identified the P300 modulation by probability density in audition and vision which is in line with this late component’s known independence from input modality^53^.

In summary, the P300 co-varied with RTs and it originated from the right parietal lobe, associating this cortical area also with the later processing stages of event probability density in temporal anticipation.

### A cerebellar signal in temporal anticipation

We observed a positive correlation between a cerebellar ERF and RT to anticipated events, which demonstrates that cerebellar activity is modulated by event probability over time. This finding is intuitively in line with the cerebellum’s well-documented role in planning and execution of motor actions^59^ and motor timing^91^. However, recently, the cerebellum was linked to other time-based, functional domains, such as attentional modulation of perception^92^ and time interval prediction^93^. This raises the question whether the cerebellum subserves a role outside of motor processes in temporal anticipation. The cerebellum’s vast connectivity to (pre-)motor^94^ and prefrontal cortices^95^ also includes reciprocal connections to posterior parietal integration areas^96^, e.g to SPL^70^, and is often seen as an indicator that the cerebellum may contribute to non-motor, cognitive processes. In light of these anatomical connections, the close temporal proximity between cerebellar and right posterior parietal signals (Fig. 8c top) observed here may indeed reflect a functional role of the cerebellum in temporal anticipation outside of motor control. This hypothesis remains to be investigated.

In conclusion, this study reveals three dynamic prediction signals, two in higher-order parietal and posterior temporal cortical regions and one in sensorimotor cortex. We also report that event-related fields over right parietal and motor areas and in the cerebellum are sensitive to event probability density after a predicted sensory cue. These neural signals demonstrate that the human brain represents the probability density function of sensory events distributed in time – a key variable underlying temporal anticipation.

## Methods

The experiments were approved by the Ethics Committee of the University Hospital Frankfurt. Written informed consent was given by all participants prior to the experiment.

### Participants

24 healthy adults (15 female), aged 21-34 years, mean age 27 years, participated in the experiment. All were right-handed and had normal or corrected-to-normal vision, reported no hearing impairment and no history of neurological disorder. Participants received € 15 per hour. One participant was excluded from the analysis of MEG data because the anatomical MRI data were corrupted and could not be retrieved.

### Experimental task and stimuli

In the MEG booth, participants performed visual and auditory blocks of trials of a Set-Go task. In the task, a SET cue was followed by a Go cue (Fig. 2a). Participants were asked to respond as quickly as possible to the Go cue with a button press (Current Designs Inc., Philadelphia, PA, USA) using their right index finger. Participants were instructed to foveate a central black fixation dot and restrict blinking to the timespan immediately following a button press. In 10 % of trials, no Go cue was presented. In these catch trials, participants were asked not to press the button. This small percentage of catch trials was added to avoid possible strong effects of event certainty towards the end of the Go-time span^38^. A small black circle around the central fixation dot was presented onscreen for 200 ms after a button press indicating the end of the trial. The intertrial interval (ITI) was defined by the onset of the small black circle and the Set cue of the following trial. The ITI was drawn randomly from a uniform distribution (range 1.4 to 2.4. s, discretized in steps of 200 ms).

### Visual stimuli

Two simultaneously presented checker boards served as both Set and Go cues. They were projected (refresh rate 60 Hz) to the back of a gray semi-translucent screen located at a fixed distance of approximately 53 cm from the participants’ eyes. Each cue was onscreen for 50 ms. The checker boards subtended approximately 6.5 × 6.5° visual angle and comprised of 5 × 5 small black and white squares of equal size. The center of the checker boards was located to the left and right of a central fixation dot at a horizonal distance of approximately 7° visual angle and a vertical distance of 0° visual angle. The black and white pattern of the checker boards was inverted between Set and Go cues.

### Auditory stimuli

White noise bursts of 50 ms length served as Set and Go cues. Each burst featured an 8 ms cosine ramp at beginning and end. The bursts were presented at approx. 60 dB SPL above hearing threshold as determined by pure tone audiometry (1 kHz, staircase procedure). All auditory stimuli were output via a RME Fireface UCX interface to a headphone amp (Lake People GT-109) and delivered diotically via a MEG-compatible tube-based system (Eartone Gold 3A 3C, Etymotic Research, Elk Grove Village, IL, USA). Visual and auditory stimuli were generated using MatLab (The MathWorks, Natick, MA, USA) and the Psychophysics Toolbox^97^ on a Fujitsy Celsius R940 computer running Windows 7 (64 bit).

### Temporal probabilities

The time between Set and Go cues, the Go-time, was a random variable, drawn from either an exponential distribution (Equation 1) with parameter *l* = 0.33 or from its left-right flipped counterpart.

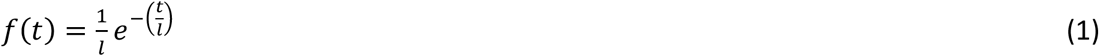

Both distributions were delayed by 0.4 s resulting in a range of Go-times from 0.4 s to 1.4 s. Sequential effects were reduced by the constraint that no more than two consecutive trials were allowed the same Go-time. The Go-time distribution was fixed for a pair of consecutive blocks of trials. Per participant the experiment consisted of four visual and four auditory blocks. A single block consisted of 200 trials of which 20 did not feature a Go cue (catch trials). Per sensory modality, in two blocks of trials, the Go-times were randomly drawn from the exponential distribution and in the other two blocks they were randomly drawn from the flipped-exponential distribution as described above.

### Models of reaction time

Hazard-rate-based and PDF-based models of RT were constructed to investigate the effect of the Go-time distribution on event anticipation. The presented exponential and flipped exponential ‘go’ time distributions are characterized by three functions, the probability density function (PDF), the cumulative distribution function (CDF), and the hazard rate (HR):

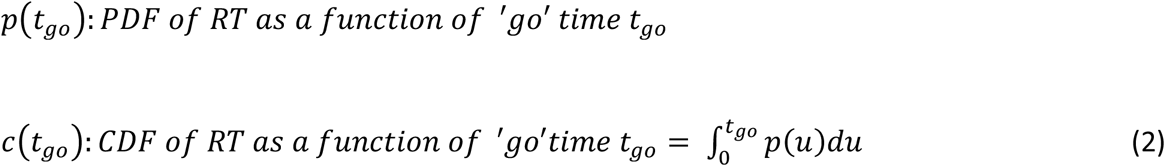

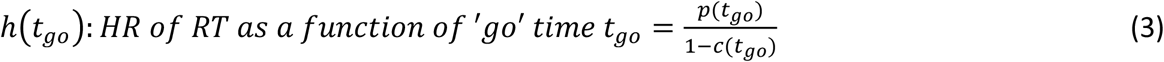

### Mirrored temporally blurred hazard rate

To arrive at the temporally blurred HR, each Go-time PDF was blurred by a Gaussian uncertainty kernel whose standard deviation linearly increases with elapsed time from a reference time point: σ = *φ* · *t*. Here, *t* is the elapsed time and *φ* is a scale factor by which the standard deviation σ of the Gaussian kernel increases. The equations for the corresponding temporally blurred functions are:

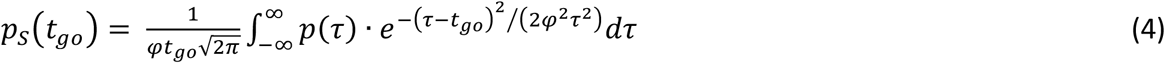

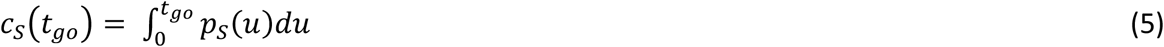

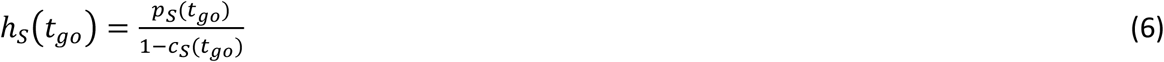

For a given Go-time *t*_*go*_ the PDF is convolved with a Gaussian kernel centered at *t*_*go*_ (Equation 4). At *t*_*go*_ = 0.4 *s* after Set cue onset the kernel has standard deviation *φ* · 0.4. Similarly at *t*_*go*_ = 1.4 *s* the kernel has standard deviation *φ* · 1.4. In the computation of the subjective PDF, the definition of the PDF was extended to the left and right of the ‘go’ time range:

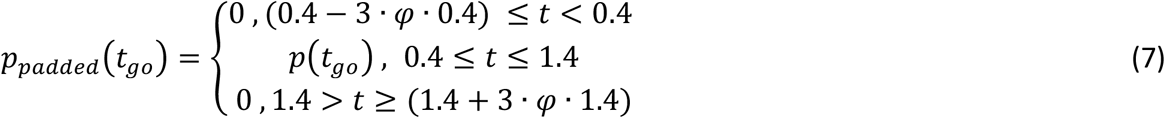

The extensions were equal to three standard deviations of the Gaussian kernel (encapsulating 99.7 % of the Gaussian uncertainty function) at the shortest and longest Go-times. Then the integral in Equation (5) was computed between these new extrema [(0.4 ™ 3 · *φ* · 0.4), (1.4 + 3 · *φ* · 1.4)] instead of the impractical interval of minus to plus infinity. The selection of *φ* = 0.21 was consistent with previous research^11,38,39,98,99^. For *φ* = 0.21 the temporal range of the extended PDF (Equation 7) becomes [0.148, 2.28] s which is also the range of integration in the computation of the subjective PDF in Equation (4). The PDF of each distribution, *p*2*t*_*go*_3 was normalized so that its integral from 0.4 s to 1.4 s was 0.9 which reflects the 10% catch trials that did not feature a Go cue. The HR was computed based on the PDF and CDF (Equation 6). To arrive at the to-be-fit HR variable, the HR was mirrored around its mean (Equation 8).

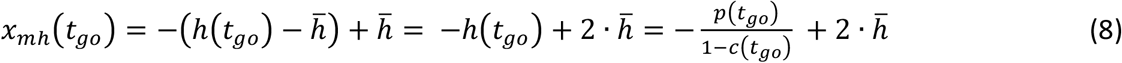

where

*x*_*mh*_ : “*mirror*” *of the hazard rate of the PDF*

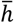: *mean HR*

### Reciprocal probabilistically blurred event probability density function

Probabilistic blurring constitutes an alternative hypothesis to the temporal blurring described above. In probabilistic blurring, the uncertainty in elapsed time estimation depends on the probability density function of event occurrence: Go-times with high probability of event occurrence are associated with low uncertainty in time estimation and vice versa, irrespective of the Go-time duration^39^. In probabilistic blurring, the standard deviation of the Gaussian kernel scales according to the PDF of event occurrence. In order to use realistic values for the standard deviation of the blurring Gaussian kernel, the minimum and maximum values were set accordingly to the temporal blurring case as

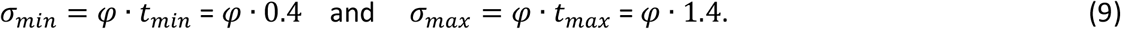

The value of *φ* was likewise set to 0.21. The PDF under investigation was then scaled so that its minimum value is σ_*min*_ and its maximum value σ_*max*_.

If *p*_*min*_ and *p*_*max*_ are the minimum and maximum values respectively of the PDF under investigation then the function used for computing the standard deviation of the Gaussian kernel based on the PDF *p*(*t*) was defined as:

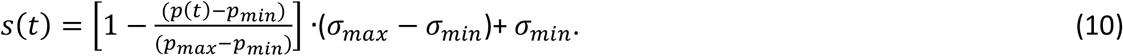

The term inside the brackets demonstrates that when the probability *p*(*t*) is low the standard deviation of the Gaussian kernel approaches σ_*max*_ while when the probability increases, *s*(*t*) approaches σ_*min*_.

Based on Equation (10) for determining the standard deviation of the Gaussian kernel the probabilistically blurred PDF *p*_*p*_(*t*) was computed as:

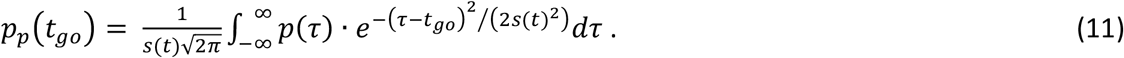

Finally, in order to implement the Gaussian blurring of Equation (11) at the extrema of Go-times, the definition of the PDF was extended to the left and right of the actual stimulus presentation interval by three standard deviations of the corresponding smoothing Gaussian kernels, similar to the temporally blurred case described in Equation (7), as:

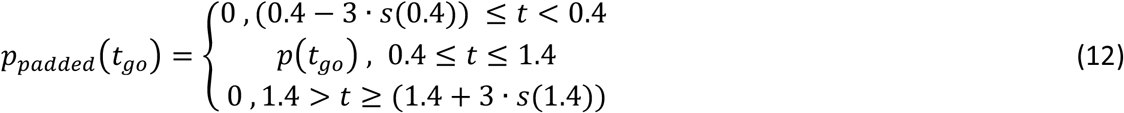

The extensions of the Go-times range depend on the standard deviation function *s*(*t*), which itself depends on the probability density function. To arrive at the to-be-fit PDF variable, the reciprocal of the PDF was computed: 1/PDF (Equation 13).

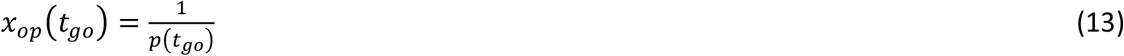

where

*x*_*op*_ : ‘*reciprocal*’ *PDF*

To illustrate the effect of the nonlinear reciprocal computation on the probabilistically blurred PDF, we compared it to the linear mirroring explained above and plotted both the probabilistically blurred PDF in its mirrored and reciprocal version in Fig. 3c.

### Modeling RT with a linear model

Both variables, the mirrored, temporally blurred HR and the probabilistically blurred 1/PDF were fit to RT data (first aggregated within Go-times within participants, then averaged across participants) using a linear model. An Ordinary Least Squares (OLS) regression was employed for the computation of the regression coefficients using the MatLab (The MathWorks, Natick MA, USA) *fit* function. Adjusted *R*^2^ was used as a measure of goodness-of-fit for comparing the models’ relation to RT.

### MEG data acquisition

Neuromagnetic activity was recorded with a 275-channel system (VSM MedTech Omega, Coquitlam, Canada) equipped with axial gradiometers distributed in a helmet across the scalp in a magnetically shielded room at the Brain Imaging Center, Frankfurt. MEG data were recorded continuously with a sampling rate of 1,200 Hz. Participants were seated in an upright position and were asked to remain still during blocks of trials. Head position relative to the MEG sensors was controlled and continually monitored during each experimental block using three position indicator coils in the anatomical fiducial locations (left and right pre-auricular points and nasion). Head position was corrected if necessary between blocks using the Fieldtrip toolbox^100^. Electro-cardiogram and vertical and horizontal electro-oculogram were also measured at 1,200 Hz to identify eye blinks and movements, and the heartbeat in analysis.

### MEG data preprocessing

Continuous data were down-sampled to 600 Hz. Data were epoched separately with respect to Set and Go cues and to the button press (−0.5 to 0.72 s). Artifactual epoches and noisy MEG channels were rejected based on visual data inspection using Fieldtrip’s visual artifact rejection routines. Based on visual inspection, trials and MEG channels that featured periods of high variance due to e.g. eyeblinks or excessive movement, during the time span of interest were discarded from the analysis. Heartbeat artifacts were removed using independent component analysis. Drifts in MEG channels were eliminated by high-pass filtering at 0.2 Hz. Muscle activity was eliminated by low-pass filtering at 110 Hz. Finally, only trials with 0.05 s < RT < 0.75 s were used in the analysis of MEG data and in modeling of RT. The trial selection process left N = 31,962 trials for analysis (3.5% of all trials removed).

### Magnetic resonance imaging data acquisition

To build participant-specific forward models for MEG source reconstruction, structural magnetic resonance imaging (MRI) scans (T1-weighted) were obtained for all 24 participants at the Brain Imaging Center, Frankfurt. MRI data were acquired on a Siemens 3T TRIO scanner with a voxel resolution of 1 × 1 × 1 mm^3^ on a 176 × 256 × 256 grid. Vitamin-E capsules were used to identify anatomic landmarks (left and right peri-auricular points and nasion).

### Time frequency analysis on sensor-space data

The time-frequency decomposition was calculated for each subject and each trial through a transformation based on multiplication in the frequency domain with a single taper (Hanning) with a 0.5 Hz resolution from 3 to 60 Hz in steps of 20 ms over t = [−0.5, 0.75] s using the FieldTrip toolbox function ft_freqanalysis with method ‘mtmconvol’. The correlation across Go-times between the spectral power and RT was performed at the single-trial level and not at the frame level as in the ERF case. This was due to the highly skewed distribution of the spectral power which results in a bias across frame average with significantly different numbers of trials (see Supplementary Methods, Suppl. Fig. 5). For each sensor, each frequency and time point the spectral power was correlated with the corresponding RT (Spearman rank correlation). At the group level, across all four experimental conditions, statistical significance was tested using a cluster-based permutation test on Spearman’s rho to identify channel-by-frequency-by-time clusters in which rho differs from zero (cluster-forming *α* = 0.05, 1000 permutations).

### Time frequency analysis on source-space data

The above correlation-based analysis on sensor-space data was also performed at the source-level representation of the time-frequency data. This source-space for each participant comprised of the individual’s cortical mantle representation, as extracted from a structural MR scan. This source space was fused with a whole-cortex parcellation atlas (AAL atlas^50^) in order to relate findings to known anatomical areas and assist the interpretation of results. Within the frequency ranges of 7–12 Hz, 15–22 Hz and 23–30 Hz and within each source, the source-level data were averaged within a time span of 100 ms, centered at t = [−0.35, −0.25, −0.15, −0.05, 0.05, 0.15, 0.25, 0.35] s relative to the Go cue. This range of time windows was selected for all four conditions based on the results of the sensor-level analysis. The inverse solution was derived by the DICS beamformer^49^. For each of the three frequency bands-of-interest, the cross-spectral density matrix used for the inverse solution was derived by averaging the cross-spectral density matrices across all time points and frequencies of interest. Then the spectral coefficients of each trial were projected through the common spatial filter from the DICS and the Spearman rank correlation between single-trial source-level spectral power and single-trial RT was computed. At the group level, across all four experimental conditions, and for each of the time windows defined above, statistical significance was tested using a cluster-based permutation test on Spearman’s rho to identify clusters of sources in which rho differs from zero (cluster-forming *α* = 0.05, 1000 permutations).

An additional correlation analysis was performed to directly investigate the relationship between the *event PDF* and source-level power. At the single-participant level and within each condition, the single-trial RTs were fit with the reciprocal probabilistically blurred event PDF variable as described above. These fits were computed in running batches of N = 15 RTs to capture potential fluctuations in average RT over the course of the two experimental blocks of a single condition. In each condition, all batches of fit PDF-based model were concatenated and Spearman correlated with source-level power at the single-trial level within each frequency band-of-interest followed by a cluster-based permutation test as described above. This analysis was performed separately for four windows of 100 ms length, covering the time span of t = [−0.4, 0 s] relative to the Go cue. Within each frequency band-of-interest, the four resultant correlation clusters were averaged across time for plotting.

### Event related fields analysis on sensor-space data

Within each condition, the single-participant MEG data was time-locked to the Go cue. The data was aggregated within adjacent pairs of consecutive Go-times (frames): For the first frame, the activity from all trials with Go-times = [0.4, 0.4167] s was averaged; the second frame comprised Go-times = [0.4333, 0.45] s and so forth. This procedure was also applied to the RT. The aggregation reduced the number of unique Go-times from 60 to 30 (= 30 frames). Before the averaging of the ERF, the data was baselined either to the pre-SET or to the pre-GO period. The pre-SET period (t = [−0.5, 0] s) was selected in the analysis of activity preceding the Go cue, whereas the pre-GO period (t = [−0.5, 0] s) was selected in the analysis of activity following the Go cue. At the single-participant level, for each condition and for each channel-by-time-point duplet, Spearman rank correlation was computed between the 30 frames of MEG data and the 30 averaged RTs. Note that the resultant rho has the same dimensionality as the within-participant grand average of MEG data (channels-by-time-points). At the group level, within each experimental condition statistical significance was tested using a cluster-based permutation test on Spearman’s rho to identify channel-by-time clusters in which rho differs from zero (cluster-forming *α* = 0.05, 1000 permutations).

### Event related fields analysis on source-space data

We used a linearly constrained minimum variance beamformer LCMV^101^. The co-variance matrix used for computation of the spatial filter was derived from the average across trials. The time period used for computation of the covariance matrix was t = [−0.5, 0.75] s relative to the Go cue. The average, baselined ERF for the computation of the covariance matrix was derived by averaging all trials from all frames so that the spatial filters are common for all frames. The sensor-level average ERF for each frame (see above) was projected to source space through the common spatial filters. For each source location, Spearman correlation was computed between the per-frame averaged ERF and the corresponding RTs. This was computed for each participant and experimental condition. Rho was averaged across time within windows-of-interest that were chosen based on the sensor-space clusters (30 ms windows centered around 0.25 and 0.4 s post Go) and one additional 30 ms window centered at 0.1 s post Go to probe for early correlations before the button response, all for each participant and experimental condition. At the group level, across all experimental conditions, the non-parametric cluster-based permutation statistics were computed on averaged Spearman’s rho to identify clusters of sources in which rho differs from zero (cluster-forming *α* = 0.05, 1000 permutations).

An additional correlation analysis was performed to directly investigate the relationship between the *event PDF* and source-level ERF. At the single-participant level and within each condition, the single-trial RTs were fit with the reciprocal probabilistically blurred event PDF variable as described above. These fits were computed in running batches of N = 15 RTs to capture potential fluctuations in average RT over the course of the two experimental blocks of a single condition. In each condition, all batches of fit PDF-based model were concatenated and Spearman correlated with the per-frame averaged ERF for each source location for each participant, using the regime described above. At the group level, across all experimental conditions, the non-parametric cluster-based permutation statistics were computed on Spearman’s rho to identify clusters of sources in which rho differs from zero (cluster-forming *α* = 0.05, 1000 permutations).

## References

1 Starkweather, C. K., Babayan, B. M., Uchida, N. & Gershman, S. J. Dopamine reward prediction errors reflect hidden-state inference across time. Nature neuroscience 20, 581–589 (2017).

2 Clark, R. E. & Squire, L. R. Classical conditioning and brain systems: the role of awareness. Science (New York, N.Y.) 280, 77–81 (1998).

3 Fiorillo, C. D., Newsome, W. T. & Schultz, W. The temporal precision of reward prediction in dopamine neurons. Nature neuroscience 11, 966–973 (2008).

4 Summerfield, C. & de Lange, F. P. Expectation in perceptual decision making: neural and computational mechanisms. Nature reviews. Neuroscience 15, 745–756 (2014).

5 Carnevale, F., de Lafuente, V., Romo, R., Barak, O. & Parga, N. Dynamic Control of Response Criterion in Premotor Cortex during Perceptual Detection under Temporal Uncertainty. Neuron 86, 1067–1077 (2015).

6 Riehle, A., Grun, S., Diesmann, M. & Aertsen, A. Spike synchronization and rate modulation differentially involved in motor cortical function. Science (New York, N.Y.) 278, 1950–1953 (1997).

7 Egger, S. W., Le, N. M. & Jazayeri, M. A neural circuit model for human sensorimotor timing. Nature communications 11, 3933 (2020).

8 Mazzucato, L., La Camera, G. & Fontanini, A. Expectation-induced modulation of metastable activity underlies faster coding of sensory stimuli. Nature neuroscience 22, 787–796 (2019).

9 Jaramillo, S. & Zador, A. M. The auditory cortex mediates the perceptual effects of acoustic temporal expectation. Nature neuroscience 14, 246–251 (2011).

10 Doherty, J. R., Rao, A., Mesulam, M. M. & Nobre, A. C. Synergistic effect of combined temporal and spatial expectations on visual attention. The Journal of neuroscience : the official journal of the Society for Neuroscience 25, 8259–8266 (2005).

11 Janssen, P. & Shadlen, M. N. A representation of the hazard rate of elapsed time in macaque area LIP. Nature neuroscience 8, 234–241 (2005).

12 Nobre, A. C. & van Ede, F. Anticipated moments: temporal structure in attention.Nature reviews. Neuroscience 19, 34–48 (2018).

13 Cravo, A. M., Rohenkohl, G., Wyart, V. & Nobre, A. C. Endogenous modulation of low frequency oscillations by temporal expectations. Journal of neurophysiology 106, 2964–2972 (2011).

14 Cui, X., Stetson, C., Montague, P. R. & Eagleman, D. M. Ready…go: Amplitude of the FMRI signal encodes expectation of cue arrival time. PLoS biology 7, e1000167 (2009).

15 de Hemptinne, C., Nozaradan, S., Duvivier, Q., Lefevre, P. & Missal, M. How do primates anticipate uncertain future events? The Journal of neuroscience : the official journal of the Society for Neuroscience 27, 4334–4341 (2007).

16 Oswal, A., Ogden, M. & Carpenter, R. H. The time course of stimulus expectation in a saccadic decision task. Journal of neurophysiology 97, 2722–2730 (2007).

17 Schoffelen, J. M., Oostenveld, R. & Fries, P. Neuronal coherence as a mechanism of effective corticospinal interaction. Science (New York, N.Y.) 308, 111–113 (2005).

18 Tsao, A., Yousefzadeh, S. A., Meck, W. H., Moser, M.-B. & Moser, E. I. The neural bases for timing of durations. Nature Reviews Neuroscience 23, 646–665 (2022).

19 Sharma, J., Sugihara, H., Katz, Y., Schummers, J., Tenenbaum, J. & Sur, M. Spatial Attention and Temporal Expectation Under Timed Uncertainty Predictably Modulate Neuronal Responses in Monkey V1. Cerebral cortex (New York, N.Y. : 1991) 25, 2894–2906 (2015).

20 Ghose, G. M. & Maunsell, J. H. Attentional modulation in visual cortex depends on task timing. Nature 419, 616–620 (2002).

21 Demarchi, G., Sanchez, G. & Weisz, N. Automatic and feature-specific prediction-related neural activity in the human auditory system. Nature communications 10, 3440 (2019).

22 Kok, P., Jehee, J. F. & de Lange, F. P. Less is more: expectation sharpens representations in the primary visual cortex. Neuron 75, 265–270 (2012).

23 van Ede, F., de Lange, F., Jensen, O. & Maris, E. Orienting attention to an upcoming tactile event involves a spatially and temporally specific modulation of sensorimotor alpha- and beta-band oscillations. The Journal of neuroscience : the official journal of the Society for Neuroscience 31, 2016–2024 (2011).

24 Kok, P. & de Lange, F. in An introduction to model-based cognitive neuroscience (eds B. U. Forstmann & E. J. Wagenmakers) 221–244 (Springer Science + Business Media, 2015).

25 Xu, S., Jiang, W., Poo, M. M. & Dan, Y. Activity recall in a visual cortical ensemble. Nature neuroscience 15, 449–455, s441-442 (2012).

26 de Lange, F. P., Heilbron, M. & Kok, P. How Do Expectations Shape Perception? Trends in cognitive sciences 22, 764–779 (2018).

27 Andersen, R. A. & Cui, H. Intention, action planning, and decision making in parietal-frontal circuits. Neuron 63, 568–583 (2009).

28 Aflalo, T. et al. Neurophysiology. Decoding motor imagery from the posterior parietal cortex of a tetraplegic human. Science (New York, N.Y.) 348, 906–910 (2015).

29 Andersen, R. A., Aflalo, T., Bashford, L., Bjånes, D. & Kellis, S. Exploring Cognition with Brain–Machine Interfaces. Annual review of psychology 73, 131–158 (2022).

30 Luce, R. D. Response Times: Their Role in Inferring Elementary Mental Organization., (Oxford Univ. Press; Clarendon Press, 1986).

31 Nobre, A., Correa, A. & Coull, J. The hazards of time. Current opinion in neurobiology 17, 465–470 (2007).

32 Bueti, D., Bahrami, B., Walsh, V. & Rees, G. Encoding of temporal probabilities in the human brain. The Journal of neuroscience : the official journal of the Society for Neuroscience 30, 4343–4352 (2010).

33 Tsunoda, Y. & Kakei, S. Reaction time changes with the hazard rate for a behaviorally relevant event when monkeys perform a delayed wrist movement task. Neuroscience letters 433, 152–157 (2008).

34 Pasquereau, B. & Turner, R. S. Dopamine neurons encode errors in predicting movement trigger occurrence. Journal of neurophysiology 113, 1110–1123 (2015).

35 Motiwala, A., Soares, S., Atallah, B. V., Paton, J. J. & Machens, C. K. Efficient coding of cognitive variables underlies dopamine response and choice behavior. Nature neuroscience 25, 738–748 (2022).

36 Barlow R E, M. A. W., Proschan F. Properties of probability distributions with monotone hazard rate. Ann Math Stat 37, 1574–1592 (1966).

37 Rice, J. & Rosenblatt, M. Estimators of the log survivor function and hazard function. Sankhya: The Indian Journal of Statistics Series A, 60-78 (1976).

38 Grabenhorst, M., Maloney, L. T., Poeppel, D. & Michalareas, G. Two sources of uncertainty independently modulate temporal expectancy. Proceedings of the National Academy of Sciences of the United States of America 118 (2021).

39 Grabenhorst, M., Michalareas, G., Maloney, L. T. & Poeppel, D. The anticipation of events in time. Nature communications 10, 5802 (2019).

40 Niemi, P. & Näätänen, R. Foreperiod and simple reaction time. Psychol. Bull. 89, 133–162 (1981).

41 Repp, B. H. & Su, Y. H. Sensorimotor synchronization: a review of recent research (2006-2012). Psychonomic bulletin & review 20, 403–452 (2013).

42 Buhusi, C. V. & Meck, W. H. What makes us tick? Functional and neural mechanisms of interval timing. Nature reviews. Neuroscience 6, 755–765 (2005).

43 Gibbon, J. Scalar expectancy theory and Weber’s law in animal timing. Psychological review 84, 279–325 (1977).

44 Gastaut, H. [Electrocorticographic study of the reactivity of rolandic rhythm]. Rev Neurol (Paris) 87, 176–182 (1952).

45 P., K. S., C., L. E., B., R. R. & J., F. J. Increases in Alpha Oscillatory Power Reflect an Active Retinotopic Mechanism for Distracter Suppression During Sustained Visuospatial Attention. Journal of neurophysiology 95, 3844–3851 (2006).

46 Deng, Y., Choi, I. & Shinn-Cunningham, B. Topographic specificity of alpha power during auditory spatial attention. NeuroImage 207, 116360 (2020).

47 Jutai, J. W., Gruzelier, J. H. & Connolly, J. F. Spectral analysis of the visual evoked potential (VEP): effects of stimulus luminance. Psychophysiology 21, 665–672 (1984).

48 Pfurtscheller, G. & Lopes da Silva, F. H. Event-related EEG/MEG synchronization and desynchronization: basic principles. Clinical neurophysiology : official journal of the International Federation of Clinical Neurophysiology 110, 1842–1857 (1999).

49 Gross, J., Kujala, J., Hämäläinen, M., Timmermann, L., Schnitzler, A. & Salmelin, R. Dynamic imaging of coherent sources: Studying neural interactions in the human brain. Proceedings of the National Academy of Sciences 98, 694 (2001).

50 Rolls, E. T., Huang, C.-C., Lin, C.-P., Feng, J. & Joliot, M. Automated anatomical labelling atlas 3. NeuroImage 206, 116189 (2020).

51 Bauer, A.-K. R., Ede, F. v., Quinn, A. J. & Nobre, A. C. Rhythmic Modulation of Visual Perception by Continuous Rhythmic Auditory Stimulation. The Journal of Neuroscience 41, 7065–7075 (2021).

52 Nobre, A. C. & van Ede, F. Attention in flux. Neuron 111, 971–986 (2023).

53 Luck, S. J. An Introduction to the Event-Related Potential Technique. Second edn, (The MIT Press, 2014).

54 Polich, J. in The Oxford Handbook of Event-Related Potential Components (eds E. S. Kappenman & S. J. Luck) (Oxford University Press, 2012).

55 Polich, J. Probability and inter-stimulus interval effects on the P300 from auditory stimuli. International Journal of Psychophysiology 10, 163–170 (1990).

56 Tueting, P., Sutton, S. & Zubin, J. Quantitative evoked potential correlates of the probability of events. Psychophysiology 7 3, 385–394 (1970).

57 Duncan-Johnson, C. C. & Donchin, E. On quantifying surprise: the variation of event-related potentials with subjective probability. Psychophysiology 14, 456–467 (1977).

58 Gross, J. et al. The neural basis of intermittent motor control in humans. Proceedings of the National Academy of Sciences of the United States of America 99, 2299–2302 (2002).

59 Allen, G. I. & Tsukahara, N. Cerebrocerebellar communication systems. Physiological Reviews 54, 957–1006 (1974).

60 Ivry, R. B. & Spencer, R. M. The neural representation of time. Current opinion in neurobiology 14, 225–232 (2004).

61 Xu, J. et al. Tractography-based Parcellation of the Human Middle Temporal Gyrus. Sci Rep 5, 18883 (2015).

62 Giraud, A. L. et al. Contributions of sensory input, auditory search and verbal comprehension to cortical activity during speech processing. Cerebral cortex (New York, N.Y. : 1991) 14, 247–255 (2004).

63 Leon, M. I. & Shadlen, M. N. Representation of time by neurons in the posterior parietal cortex of the macaque. Neuron 38, 317–327 (2003).

64 Battelli, L. et al. Unilateral right parietal damage leads to bilateral deficit for high-level motion. Neuron 32, 985–995 (2001).

65 Husain, M., Shapiro, K., Martin, J. & Kennard, C. Abnormal temporal dynamics of visual attention in spatial neglect patients. Nature 385, 154–156 (1997).

66 Agosta, S. et al. The Pivotal Role of the Right Parietal Lobe in Temporal Attention. Journal of cognitive neuroscience 29, 805–815 (2017).

67 Battelli, L., Cavanagh, P. & Thornton, I. M. Perception of biological motion in parietal patients. Neuropsychologia 41, 1808–1816 (2003).

68 Hayashi, M. J. & Ivry, R. B. Duration Selectivity in Right Parietal Cortex Reflects the Subjective Experience of Time. The Journal of neuroscience : the official journal of the Society for Neuroscience 40, 7749–7758 (2020).

69 Rolls, E. T., Deco, G., Huang, C. C. & Feng, J. The human posterior parietal cortex: effective connectome, and its relation to function. Cerebral cortex (New York, N.Y. : 1991) 33, 3142–3170 (2023).

70 Wang, J. et al. Convergent functional architecture of the superior parietal lobule unraveled with multimodal neuroimaging approaches. Human brain mapping 36, 238–257 (2015).

71 Wang, J. et al. Functional topography of the right inferior parietal lobule structured by anatomical connectivity profiles. Human brain mapping 37, 4316–4332 (2016).

72 Zhang, C. Y. et al. Partially Mixed Selectivity in Human Posterior Parietal Association Cortex. Neuron 95, 697–708.e694 (2017).

73 Howard, C. J., Boulton, H., Bedwell, S. A., Boatman, C. A., Roberts, K. L. & Mitra, S. Low-Frequency Repetitive Transcranial Magnetic Stimulation to Right Parietal Cortex Disrupts Perception of Briefly Presented Stimuli. Perception 48, 346–355 (2019).

74 Bareham, C. A., Georgieva, S. D., Kamke, M. R., Lloyd, D., Bekinschtein, T. A. & Mattingley, J. B. Role of the right inferior parietal cortex in auditory selective attention: An rTMS study. Cortex 99, 30–38 (2018).

75 Battelli, L., Walsh, V., Pascual-Leone, A. & Cavanagh, P. The ‘when’ parietal pathway explored by lesion studies. Current opinion in neurobiology 18, 120–126 (2008).

76 Corbetta, M. & Shulman, G. L. Control of goal-directed and stimulus-driven attention in the brain. Nature reviews. Neuroscience 3, 201–215 (2002).

77 Andersen, R. A. & Buneo, C. A. Intentional maps in posterior parietal cortex. Annual review of neuroscience 25, 189–220 (2002).

78 van Dijk, H., Schoffelen, J. M., Oostenveld, R. & Jensen, O. Prestimulus oscillatory activity in the alpha band predicts visual discrimination ability. The Journal of neuroscience : the official journal of the Society for Neuroscience 28, 1816–1823 (2008).

79 Hanslmayr, S., Aslan, A., Staudigl, T., Klimesch, W., Herrmann, C. S. & Bäuml, K. H. Prestimulus oscillations predict visual perception performance between and within subjects. NeuroImage 37, 1465–1473 (2007).

80 Klimesch, W., Sauseng, P. & Hanslmayr, S. EEG alpha oscillations: the inhibition-timing hypothesis. Brain Res Rev 53, 63–88 (2007).

81 Klimesch, W. α-band oscillations, attention, and controlled access to stored information. Trends in cognitive sciences 16, 606–617 (2012).

82 Misselhorn, J., Friese, U. & Engel, A. K. Frontal and parietal alpha oscillations reflect attentional modulation of cross-modal matching. Scientific Reports 9, 5030 (2019).

83 Worden, M. S., Foxe, J. J., Wang, N. & Simpson, G. V. Anticipatory biasing of visuospatial attention indexed by retinotopically specific alpha-band electroencephalography increases over occipital cortex. The Journal of neuroscience : the official journal of the Society for Neuroscience 20, Rc63 (2000).

84 Sauseng, P. et al. A shift of visual spatial attention is selectively associated with human EEG alpha activity. The European journal of neuroscience 22, 2917–2926 (2005).

85 Nieuwenhuis, S., Aston-Jones, G. & Cohen, J. D. Decision making, the P3, and the locus coeruleus-norepinephrine system. Psychological bulletin 131, 510–532 (2005).

86 O’Connell, R. G. & Kelly, S. P. Neurophysiology of Human Perceptual Decision-Making. Annual review of neuroscience 44, 495–516 (2021).

87 Dehaene, S., Changeux, J. P., Naccache, L., Sackur, J. & Sergent, C. Conscious, preconscious, and subliminal processing: a testable taxonomy. Trends in cognitive sciences 10, 204–211 (2006).

88 Polich, J. Updating P300: an integrative theory of P3a and P3b. Clinical neurophysiology : official journal of the International Federation of Clinical Neurophysiology 118, 2128–2148 (2007).

89 Squires, N. K., Squires, K. C. & Hillyard, S. A. Two varieties of long-latency positive waves evoked by unpredictable auditory stimuli in man. Electroencephalogr Clin Neurophysiol 38, 387–401 (1975).

90 Donchin, E. Presidential address, 1980. Surprise!…Surprise? Psychophysiology 18, 493–513 (1981).

91 Ivry, R. B. & Keele, S. W. Timing Functions of The Cerebellum. Journal of cognitive neuroscience 1, 136–152 (1989).

92 Breska, A. & Ivry, R. B. The human cerebellum is essential for modulating perceptual sensitivity based on temporal expectations. eLife 10 (2021).

93 Breska, A. & Ivry, R. B. Double dissociation of single-interval and rhythmic temporal prediction in cerebellar degeneration and Parkinson’s disease. Proceedings of the National Academy of Sciences of the United States of America 115, 12283–12288 (2018).

94 Middleton, F. A. & Strick, P. L. Basal ganglia and cerebellar loops: motor and cognitive circuits. Brain Res Brain Res Rev 31, 236–250 (2000).

95 Ramnani, N. The primate cortico-cerebellar system: anatomy and function. Nature reviews. Neuroscience 7, 511–522 (2006).

96 Bostan, A. C., Dum, R. P. & Strick, P. L. Cerebellar networks with the cerebral cortex and basal ganglia. Trends in cognitive sciences 17, 241–254 (2013).

97 Brainard, D. H. The Psychophysics Toolbox. Spatial vision 10, 433–436 (1997).

98 Gibbon, J., Malapani, C., Dale, C. L. & Gallistel, C. Toward a neurobiology of temporal cognition: advances and challenges. Current opinion in neurobiology 7, 170–184 (1997).

99 Rakitin, B. C., Gibbon, J., Penney, T. B., Malapani, C., Hinton, S. C. & Meck, W. H. Scalar expectancy theory and peak-interval timing in humans. Journal of experimental psychology. Animal behavior processes 24, 15–33 (1998).

100 Oostenveld, R., Fries, P., Maris, E. & Schoffelen, J. M. FieldTrip: Open source software for advanced analysis of MEG, EEG, and invasive electrophysiological data. Comput Intell Neurosci 2011, 156869 (2011).

101 Veen, B. D. V., Drongelen, W. V., Yuchtman, M. & Suzuki, A. Localization of brain electrical activity via linearly constrained minimum variance spatial filtering. IEEE Transactions on Biomedical Engineering 44, 867–880 (1997).

