## Supplementary Material for "Neural signatures of temporal anticipation in human cortex represent event probability density"

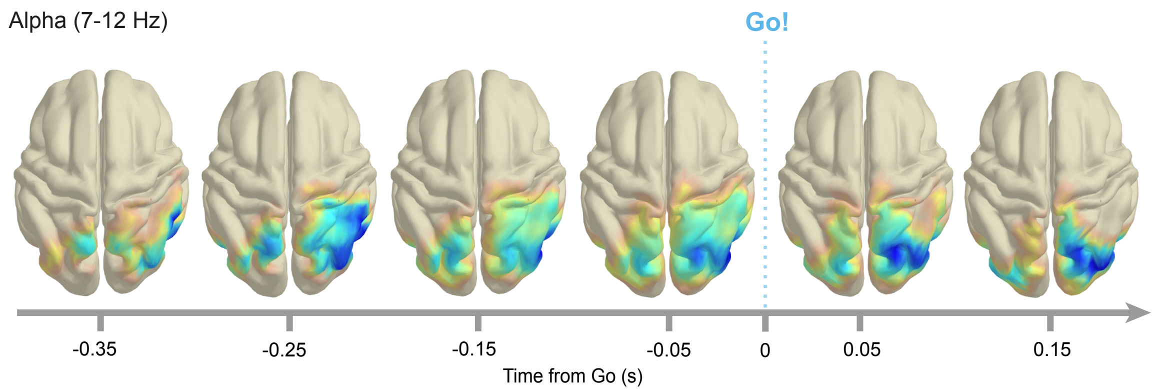


**Supplementary Fig. 1 | Right lateralization of source-level correlation clusters in the alpha band.** Spearman correlation computed between single-trial RT and single-trial alpha power averaged across frequencies (7-12 Hz). Plots use an individual colormap for each time window (see Fig. S2).

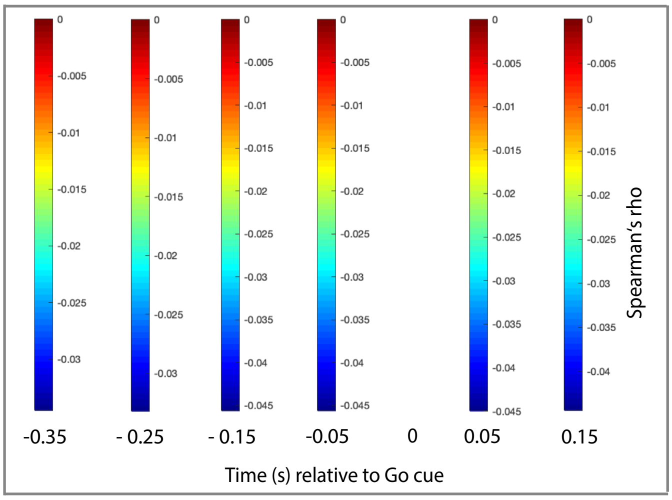


**Supplementary Fig. 2 | Colorbars depict Spearman's rho over time**. Individual colorbars for re-plotting of correlation (source-level alpha power and RT) shown in Fig. 6 (top row) and in Fig. S1.

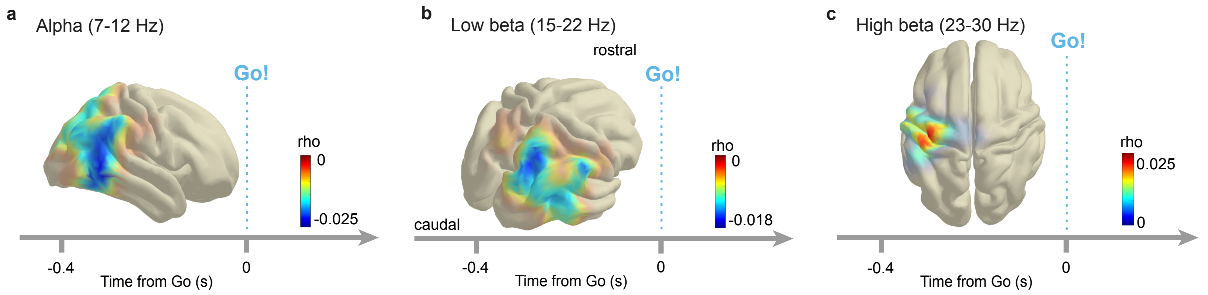


**Supplementary Fig. 3 | Event PDF is represented in cortical key regions prior to the anticipated sensory cue. a.** Cortical topography of across-condition Spearman correlation clusters (alpha (7–12 Hz) power, fit PDF-based model of RT), averaged across time (-0.4 to 0 s, P = [0.001, 0.006, 0.009, 0.01], Methods). Negative correlation covers inferior parietal lobule and the posterior middle temporal gyrus area. **b.** Across-condition Spearman correlation clusters (low beta (15–22 Hz) power, fit PDF-based model of RT), averaged across time (-0.4 to 0 s, P = [0.0035, 0.0085, 0.019, 0.021]). Negative correlation cluster in right superior parietal lobule (SPL). **c.** Across-condition Spearman correlation clusters (high beta (23–30 Hz) power, fit PDF-based model of RT), averaged across time (-0.4 to 0 s, P = [0.18, 0.012, 0.013, 0.001]). Positive correlation clusters in left sensorimotor cortex prior to Go cue. P values uncorrected.

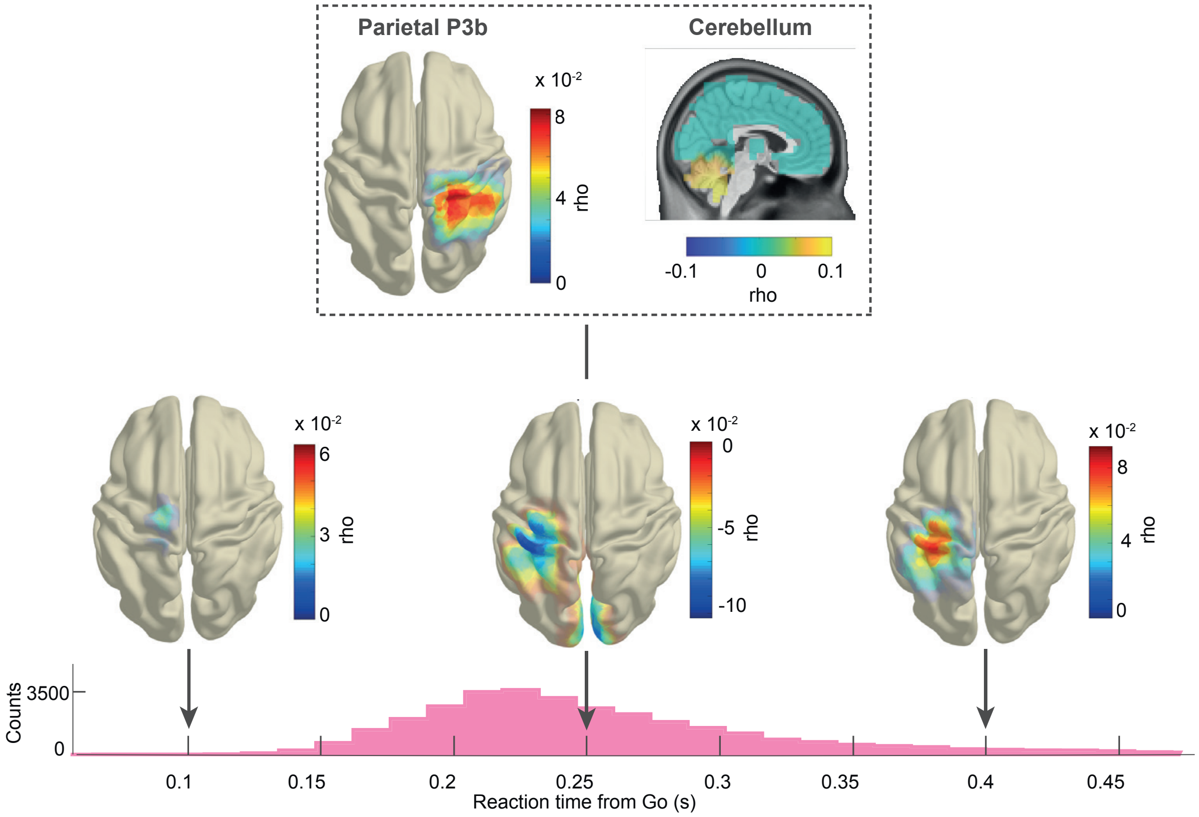


**Supplementary Fig. 4 | Parietal, cerebellar and sensorimotor event-related fields correlate with fit PDF-based model of RT.** Topography of across-condition source-level Spearman correlation clusters (ERF, PDF-based model of RT). At 0.25 s post-Go cue, positive correlation clusters in right parietal cortex (P3b, P = 0.024) and cerebellum (P = 0.006, both top) and a negative cluster in left sensorimotor cortex (P = 0.002, bottom). At 0.1 s, the cluster test identified a small number of sources in medial motor cortex but the cluster was not significant (P = 0.70). At 0.4 s, positive correlation clusters in left sensorimotor cortex (P = 0.002). Bottom: RT histogram from Fig. 2c for orientation.

### Supplementary Methods

**A note on correlation bias.** In the ERF analysis, trials were aggregated in 30 frames of equal duration with respect to Go-time. The number of trials in each frame varied considerably, as the high probability frames contained many trials, whereas the frames with low event probability contained only few trials. Within each frame, the RT and the MEG data were respectively averaged for each subject, condition, channel and time point. So there is an obvious question whether averaging a different number of trials within each frame introduces a bias in the correlation between RT and MEG data.

We investigated this possible confound with simulated data resembling the distributions in the actual MEG data. The evoked MEG data had a distribution close to a Gaussian centered around zero while the TFR data resembled a highly skewed Weibul distribution with a long exponential tail. We simulated random data with similar distributions. RT was simulated by the 1/PDF linear model that was presented in the initial behavioral analysis.

The Go-time frames had the same distribution of trials as in the actual data. The simulated MEG data was then averaged within frames and the 30 frame values were correlated with the 30 RT simulated values. As the simulated MEG data was random, also the correlations with simulated RT values was expected to be around zero with no monotonic structure. Any monotonic structure would hint towards an artifact from the averaging of a different number of trials per frame, as the trial number decreases or increases monotonically, depending on the event distribution (exponential: decreasing, flipped-exponential: increasing).

This procedure was repeated 10000 times with each time having new random simulated MEG data. The histogram of the resultant 10000 correlation values was plotted for the simulated TFR cases (Suppl. Fig. 5a) and ERF cases (Suppl. Fig. 5b) .

In the ERF case, no bias was observed in the distribution of rho, as it was centered round zero. In contrast, a significant bias was observed in the TFR case. In this case the distribution of rho was shifted away from zero towards negative values which should not be the case as the simulated MEG data was random. Clearly this is an artifact of averaging a different number of trials per frame. The bias seems to be dominant in the frames with the small number of trials, where the average tends to assume higher values, as compared to frames with many trials. The reason for this offset is the highly skewed distribution with some trials having very large values. Such trials have a dominating effect in frames with few trials and a much smaller effect in frames with many trials. That is why the bias seems to monotonically increase as one progresses from high probability frames to the less probable ones with fewer and fewer trials. Based on these observations we decided in the actual data to follow different strategies for studying the correlation of the RT with the MEG ERF and the TFR data sets.

For the ERF data, as no bias seems to be present due to the averaging within frames, we proceeded and correlated the within-frames averaged, time-locked MEG data with the within-frames averages of RT.

For the TFR data, we correlated the data at the single-trial level, avoiding the monotonic bias in the average within bins.

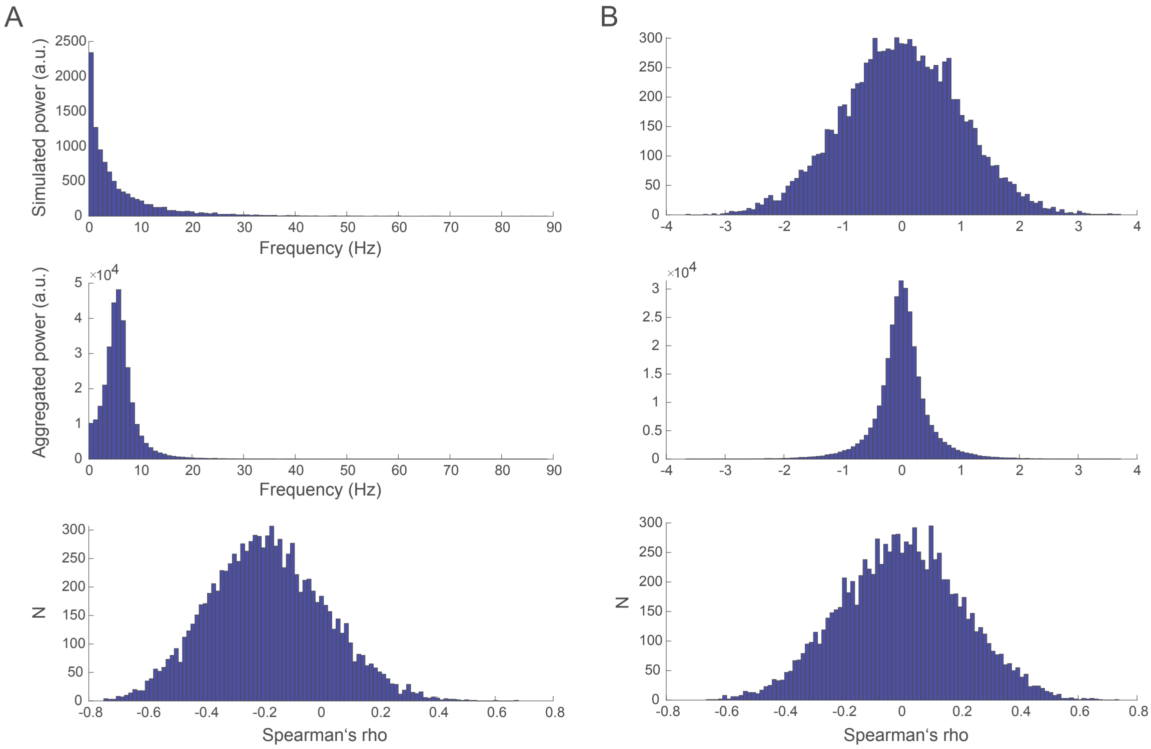


**Supplementary Fig. 5** Data simulation. **a**) Simulated MEG power (arbitrary units) (top), aggregated within frames (arbitrary units) (middle) Spearman's rho computed on aggregated simulated power and 1/PDF (bottom). **b**) Simulated ERF data (arbitrary units) (top), aggregated within frames (arbitrary units) (middle), Spearman's rho computed on aggregated simulated ERF data and 1/PDF (bottom).
